# Transcriptional activity is shaped by the chromatin landscapes in Arabidopsis

**DOI:** 10.1101/2022.06.02.494419

**Authors:** Bhagyshree Jamge, Zdravko J. Lorković, Elin Axelsson, Ramesh Yelagandula, Svetlana Akimcheva, Frédéric Berger

## Abstract

How histone variants and histone modifications shape nucleosome-mediated transcriptional repression, and how transcriptional activity shapes the enrichment of histone modifications and variants remain unclear. To clarify these relationships, we examined chromatin organization in the *Arabidopsis thaliana* genome, identifying a limited number of chromatin landscapes that distinguish transposon families and distinct groups of genes based on their transcriptional regulation. Unexpectedly, H2A variants are strong determinants of the landscape architecture. Combinations of H2A.W and four histone modifications define six domains that are occupied by specific transposon families and organized concentrically around the centromere. Moreover, H2A.Z defines transcriptional gene repression in specific domains. Expressed genes occupy four chromatin landscapes with specific RNA Polymerase II profiles. Although the composition of each chromatin landscape is invariant, they cover genes with a wide range of expression levels. Therefore, chromatin landscapes control the range of transcriptional activity, but transcriptional activity has little effect on chromatin composition.

**One Sentence Summary:** Histone variants and histone modifications build a limited number of distinct chromatin landscapes that instruct the transcriptional regulation of genes and transposons in Arabidopsis.

## Introduction

Eukaryotic genomes are packaged into chromatin, a structure defined by repeating units of ∼147 bp of DNA wrapped around a protein complex known as the nucleosome (Luger et al. 1997). Nucleosomes are essential for genome organization and function in packaging DNA (Macadangdang et al. 2014), transcription (Allis et al. 1980; Costanzi and Pehrson 1998; Giaimo et al. 2019; Meneghini, Wu, and Madhani 2003; Yelagandula et al. 2014), and genome maintenance (Rogakou et al. 1998). Notably, nucleosomes contain two copies each of histones H2A, H2B, H3, and H4. The histones associated with arrays of nucleosomes, which wrap around genes and transposons, are subject to a range of post-translational modifications (Talbert and Henikoff 2021; Tessarz and Kouzarides 2014), and chromatin states can be defined based on the characteristic combinations of histone modifications they exhibit (Ernst and Kellis 2012, 2017). Indeed, chromatin states distinguish the major domains of chromatin in the genome, comprising euchromatin (with active genes), facultative heterochromatin (with repressed genes) and constitutive heterochromatin (with transposons and repeats) (Roudier et al. 2011). They also highlight specific features such as promoters and enhancers (Ernst and Kellis 2010). How chromatin states assemble at different scales of genome organization and whether transcriptional activity shapes chromatin states, or chromatin states shape the transcriptional regulation of genes and transposons, remain unclear.

Histone variants have arisen from divergence—both within and between organisms—of their intrinsically disordered loop and tail regions, with dozens of variants in the H2A, H2B, and H3 families identified (Talbert and Henikoff 2021). All multicellular eukaryotes possess a centromeric H3 (CENPA), a DNA replication–dependent H3.1 (which is incorporated in the S phase), and a DNA replication-independent H3.3 as well as H2A.Z, H2A.X, and a DNA replication–dependent H2A variant. These variants can have important functions in chromatin state establishment. For example, nucleosomes cause RNA Polymerase II (Pol II) to stall, and the transcriptional complex must overcome the energy barrier imposed by nucleosomes to initiate transcription (Bintu et al. 2012; Bondarenko et al. 2006; Kireeva et al. 2005). This energy barrier is reduced when the first nucleosome encountered by RNA Pol II contains the histone variant H2A.Z, highlighting the potential for variants to influence transcription (Rudnizky et al. 2016; Subramanian, Fields, and Boyer 2015; Weber and Henikoff 2014).

This idea is further supported by the following observation. In fission yeast, H2A.Z participates in the initiation and termination of transcription (Albert et al. 2007; Raisner et al. 2005; Venters et al. 2011). In plants, opposing roles for H3.3 and H2A.Z have been suggested (Wollmann et al. 2017). In addition, unique types of H2A variants that are associated with transcriptional repression have evolved in animals and plants (Loppin and Berger 2020; Talbert and Henikoff 2021). In mammals, the so-called macroH2A variants have a C-terminal macro domain attached to the core by a linker enriched in lysine residues (Douet et al. 2017; Hsu et al. 2021; Sun and Bernstein 2019). In vascular plants, the H2A.W variants are further distinguished by a C-terminal KSPK motif (Lei and Berger 2020; Schmücker et al. 2021; Yelagandula et al. 2014) which favors heterochromatic silencing by directly altering H2A.W interaction with DNA (Osakabe et al. 2018; Schmücker et al. 2021; Yelagandula et al. 2014). Furthermore, a diversity of H2B variants was recently recognized in both plants (Jiang et al. 2020) and mammals (Raman et al. 2022), although the roles of these variants in transcription remains unknown. Histone variants thus represent complex, diverse nucleosome components that could regulate and organize the functional activities of chromatin.

Applying Hidden Markoff Model [ChromHMM] (Ernst and Kellis 2012, 2017), to genomic profiles of histone modifications revealed chromatin states that reflect the combination of post-transcriptional histone modifications present in mammals, Drosophila, Arabidopsis, and rice (Ernst and Kellis 2010, 2012, 2017; Kharchenko et al. 2011; Liu et al. 2018; Roudier et al. 2011; Sequeira-Mendes et al. 2014). Conserved chromatin states include H3 and H4 acetylation, which is deposited by the preinitiation complex (PIC) associated with the SAGA complex and the general transcription factor TFIID (Watanabe and Kokubo 2017). Acetylated histones assist in the docking of the chromatin remodeler SWI2/SNF2-related 1 (SWR1), which deposits H2A.Z at the transcription start sites (TSS) of expressed genes (Aslam et al. 2019). Trimethylation of lysine 4 of H3 (H3K4me3) deposited by the complex COMPASS is also correlated with the PIC, but its direct roles remain unclear (Fromm and Avramova 2014). After the initiation of transcription, RNA Pol II begins the early elongation process, which involves pausing sites (Nielsen et al. 2019). These sites are marked by the histone tail acetylation levels, H3K4me3, and H3K36me3 which are positively correlated with transcript levels (Gates, Foulds, and O’Malley 2017). Active elongation is marked by monoubiquitination of H2B (H2Bub), which requires specific ubiquitin ligases: HUB1 and Hub2 in Arabidopsis (Feng and Shen 2014) and UBE2A:RNF20/40 in human (Kim et al. 2009). This modification is associated with transcribed regions in mammals (Minsky et al. 2008) and plants (Roudier et al. 2011). With the exception of H3K9me2, which marks the 3′ ends of genes in mammals but not plants (Gates, Foulds, and O’Malley 2017; Leng et al. 2020), the histone marks or variants that control the termination of transcriptional elongation are not understood.

Compared with the wealth of data regarding the relationships of histone modifications with transcription, the role of histone variants in this process remains to be explored. In Arabidopsis, there is a clear correlation between transcript levels and the enrichment of the variant H3.3 (Stroud et al. 2012; Wollmann et al. 2012, 2017) and replicative H2A and H2A.X (Yelagandula et al. 2014)(Yelagandula et al. 2014) on gene bodies marked by H3K36me2 (Leng et al. 2020). By contrast, H2A.Z is enriched on the bodies of genes marked by H3K27me3 (Carter et al. 2018), and the histone variant H2A.W is present on transposons and repeats marked by H3K9me2 (Osakabe et al. 2021; Yelagandula et al. 2014). The complex, intimate associations between histone variants and modifications have not been fully characterized in plants or mammals. In addition, it remains unclear to what extent transcriptional activity generates these associations or whether feedback loops generate a dialog between the transcription and deposition of chromatin elements that shape chromatin organization in the genome.

Here, we analyzed how all 13 histone variants expressed in vegetative tissues associate with the 12 prominent chromatin modifications to form chromatin states in the model flowering plant *Arabidopsis thaliana.* Our findings indicate that H2A variants are major factors that differentiate among euchromatin, facultative heterochromatin, and constitutive heterochromatin. Unsupervised clustering of chromatin states showed that chromatin is organized into stereotypical Chromatin Landscapes (CLs) that distinguish distinct classes of transposons and genes with different modes of transcriptional regulation. Each CL was associated with a specific mode of transcriptional repression or activity. Remarkably, the levels of transcription did not affect the organization of the CLs, suggesting that CLs define specific modes of transcription, whereas transcription has little influence on the composition of the CL.

## Results

### Co-occurrence of H3 modifications and histone variants in nucleosomes

In Arabidopsis, homotypic nucleosomes containing a single type of H2A variant are prevalent (Osakabe et al. 2018). However, whether H3 variants also assemble in a homotypic manner and whether H2A variants preferably assemble with a specific H3 variant or histone modification have been unclear. To answer these questions, we immunoprecipitated mononucleosomes (**Fig S1A–C**) from transgenic Arabidopsis lines expressing HA-tagged H3.1 and H3.3 under the control of their native promoters (Jiang and Berger, 2017). Because tagged H3 ran slower than endogenous H3 on SDS-PAGE (**Fig S1B**), this approach allowed us to analyze the composition of H3.1 and H3.3 nucleosomes by mass spectrometry (MS) and immunoblotting. MS analysis revealed that ∼40% of nucleosomes contained both H3.1 and H3.3 (**Fig 1A**). Neither H3.1 nor H3.3 was preferentially associated with a specific H2A variant (**Fig 1B**). This was confirmed by MS analysis of nucleosomes pulled down with H2A variant-specific antibodies (**Fig S1D**). Therefore, H3 variants do not necessarily form homotypic nucleosomes, despite the presence of H3 variant-specific deposition mechanisms (Nie et al. 2014), and they do not associate with specific H2A variants.

**Fig. 1:**
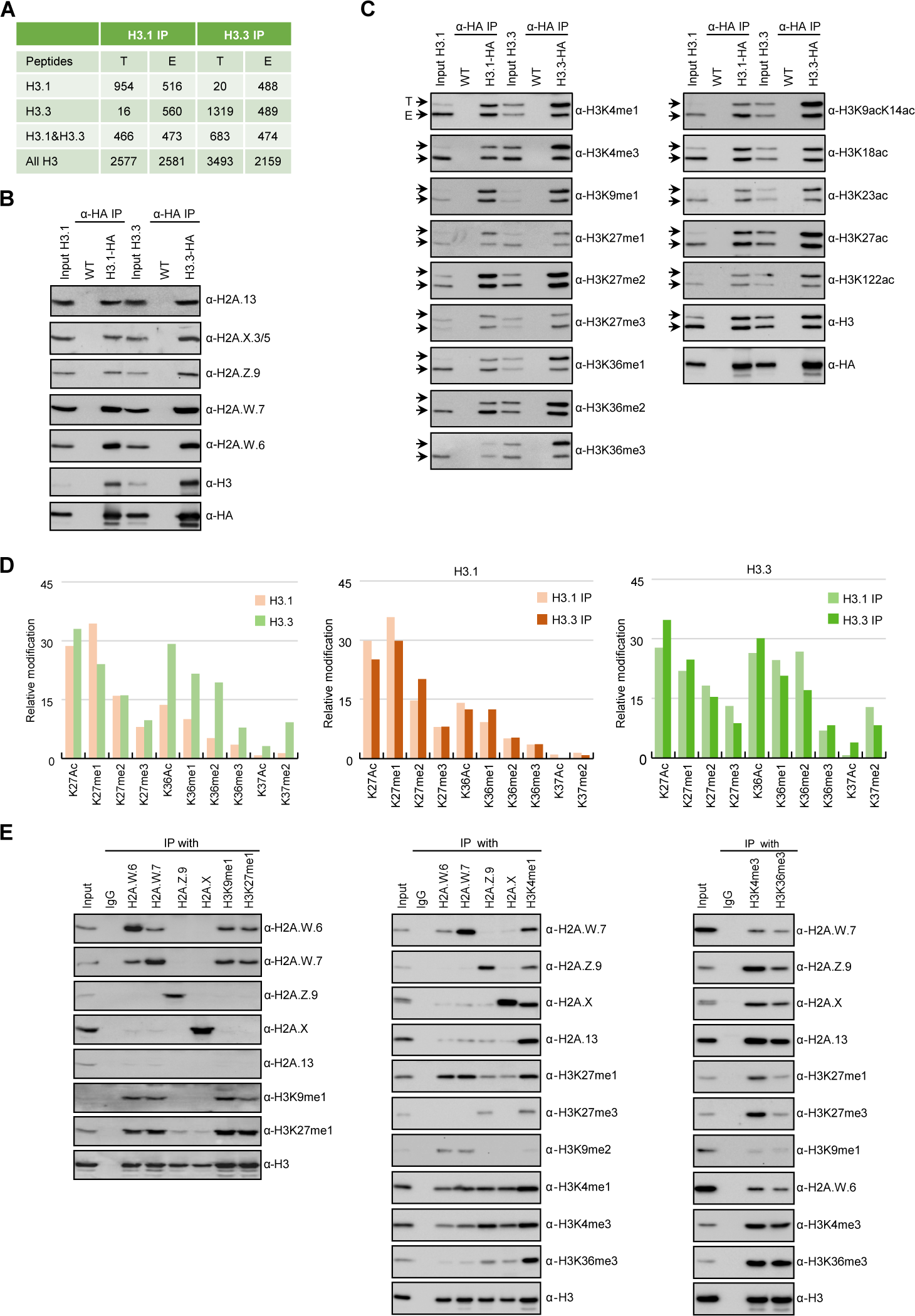
Biochemical analysis of the association between histone variants and histone marks. (A) Histone H3.1 and H3.3 form homotypic and heterotypic nucleosomes. Spectral counts of H3.1- and H3.3-specific peptides in the respective immunoprecipitates (T – transgenic, E – endogenous H31. and H3.3). (B) H2A variants do not preferentially associate with H3.1- or H3.3-containing nucleosomes. HA-tagged H3.1 and H3.3 were immunoprecipitated with HA agarose and analyzed for the presence of H2A variants by immunoblotting. (C) Histone H3 marks are present on both H3.1 and H3.3. HA-tagged H3.1 and H3.3 were immunoprecipitated with HA agarose and analyzed for the presence of H3 marks by immunoblotting. Arrows indicate transgenic (T) and endogenous (E) H3. (D) MS analysis of cumulative H3.1 and H3.3 K27 and K36 modifications in H3.1 and H3.3 immunoprecipitates (left panel). Middle and right panels represent H3.1 and H3.3 K27 and K36 modifications, respectively, as analyzed separately in H3.1 and H3.3 immunoprecipitates. (E) Co-occurrence of H2A variants and H3 marks. Mononucleosomes were immunoprecipitated with the indicated antibodies and analyzed for the presence of H2A variants and H3 marks by immunoblotting.

We also explored the degree of co-occurrence of histone modifications with H3.1 and H3.3 variants. H3.1 and H3.3 bore the same sets and comparable levels of methylation and acetylation, except that H3.3 was more strongly associated with H3 acetylation and H3K36 methylation and H3.1 with H3K27me1 (**Fig 1C**). We also analyzed H3.1 and H3.3 modifications by MS using information from both tagged H3 and endogenous H3 in the respective immunoprecipitates. MS unambiguously distinguishes H3K27 methylation and H3K36 methylation in H3.1 and H3.3 variants (**Fig S1E**). Here, we focused on H3K27 and H3K36 modifications because they are diagnostic of constitutive (H3K27me1) versus facultative (H3K27me3, silenced by Polycomb repressive complex 2 [PRC2]) heterochromatin and actively transcribed (H3K36me) regions of the genome, respectively. The levels of H3K27 and H3K36 modifications on H3.1 and H3.3 displayed the same trends regardless of whether they were precipitated with H3.1 or H3.3 nucleosomes; the same trend was also observed by comparing endogenous and transgene-encoded H3.1 and H3.3 (**Fig 1D** and **S1E–J**). As reported previously (Johnson et al., 2004), we observed some preferential association of H3.1 with H3K27me1 and H3.3 with H3K36 methylation, but this was not a strict association (**Fig 1D** and **S1E–J**). Thus, based on two different approaches, we conclude that there is no tight association between H3 modifications and the two H3 variants.

In contrast to H3 variants, homotypic nucleosomes containing either H2A.W, H2A.Z, H2A, or H2A.X showed marked enrichment of specific histone modifications. H2A.W variants were strongly associated with histone modifications of constitutive heterochromatin (H3K9me1, H3K9me2, and H3K27me1) (**Fig 1E**). The modification H3K27me3, which is associated with facultative heterochromatin, was detected only in H2A.Z nucleosomes, and H3K36me3 (marking euchromatin) was primarily detected in H2A, H2A.X, and H2A.Z nucleosomes (**Fig 1E)**. Other modifications (such as H3K4me1 and H3K4me3) displayed complex patterns of association with H2A variants and associations among themselves. Therefore, H2A but not H3 variants form homotypic nucleosomes that preferentially carry specific histone modifications that are associated with the transcriptional status of protein-coding genes or transposons.

### Identification of chromatin states using histone variants in Arabidopsis

We performed ChIP-seq using Arabidopsis seedlings to identify the combinations of all histone variants present in somatic cells and their associations with histone modifications. We included the most abundant isoforms of H2A.W (H2A.W.6 and H2A.W.7), H2A.Z (H2A.Z9 and H2A.Z.11), and replicative H2A (H2A.2 and H2A.13) in our analysis. The algorithm ChromHMM (Ernst and Kellis 2017) was used to describe chromatin states in Arabidopsis (see Methods for details). We chose to analyze the most parsimonious model based on 26 chromatin states (**Fig S2A**). Chromatin states were clustered based on the emission probability for each modification and histone variants (**Fig 2A**) and showed distinct enrichment of various marks and occupied different proportions in the genome (**Fig 2A, 2B**). For each chromatin state, we measured the average level of transcriptional activity (**Fig 2C**), the degree of enrichment in CG methylation (**Fig 2D, S2C** and **S2D**), the level of accessibility by DNase I-seq (**Fig 2E**) and the relative abundance of transposons, repeats, and elements of protein-coding genes (**Fig 2F**). Altogether, these parameters defined four groups: constitutive heterochromatin (H), facultative heterochromatin (F), euchromatin (E), and intergenic regions (I).

**Fig. 2:**
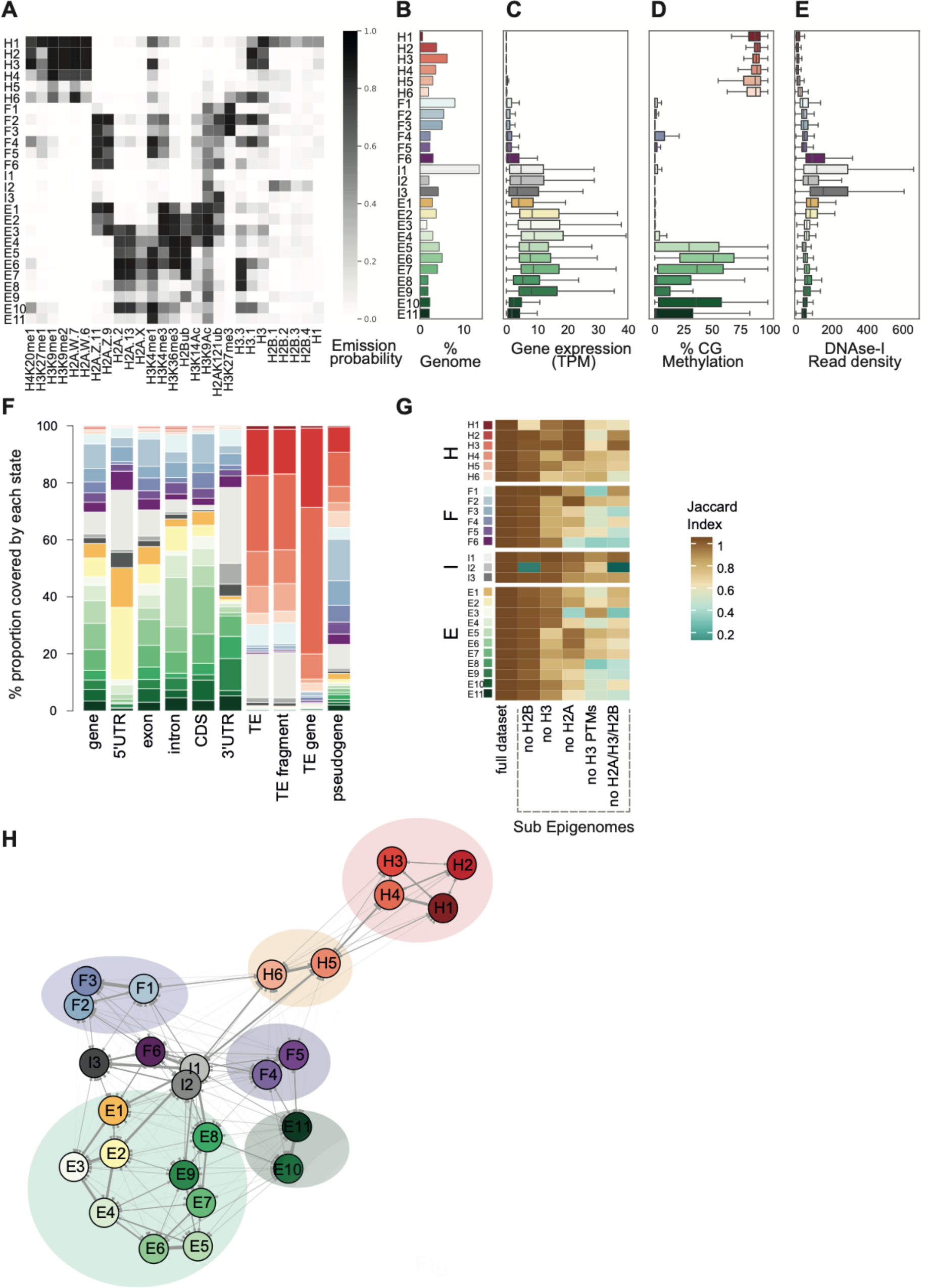
Histone variants define chromatin states in *Arabidopsis thaliana*. (A) Heatmap showing the emission probabilities for each mark/variant across the 26 chromatin states. (B) Bar plot representing the proportion of the genome occupied by each state (% Genome). (C) Boxplot showing the expression of protein-coding genes overlapping with each chromatin state in Transcripts per Million (TPM). (D) Boxplot showing CG methylation levels across chromatin states. (E) Boxplot comparing DNase I-seq read coverage across states. (F) Bar plot showing the overlap between annotated genomic features and chromatin states. (G) Heatmap showing the Jaccard similarity index between the states generated using the whole model and states using a subset of marks, i.e. excluding a set of marks the X-axis from left to right (complete data, no H2B variants, no H3 variants, no H2A variants, no histone modifications, and no variants (H2A/H3/H2B)). (H) Network diagram representing the transition matrix of all chromatin states. The thickness of the arrows is proportional to the probability of the state transition. [The thicker the connection between the states, the more likely they are nearby on a linear scale and transition from one state to other.] (The probability for self-transition in the same state was omitted from the plots to simplify the diagram.)

States H1–H6 shared the highest levels of CG methylation, with the near absence of transcriptional activity, and was primarily occupied by transposons and repeats, thus representing constitutive heterochromatin. States F1–F6 were associated with low transcriptional activity and were present over genes and half of pseudogenes, indicating that they represented facultative heterochromatin. States E1–E11 occupied expressed genes and thus comprised euchromatin. Three states (I1–I3) occupying 15% of the genome comprised non-coding regions such as transposons, repeats in regions dense with protein-coding genes, and untranslated regions of genes (3′UTR and 5′UTR). Hence, I1–I3 were classified as intergenic states. As defined by state occupancy, constitutive and facultative heterochromatin occupied 17% and 20% of the genome (**Fig S2F**), respectively, which corresponded to previous estimates (Roudier et al. 2011). Chromatin accessibility was (to some degree) inversely correlated with the degree of nucleosome density determined by MNase-seq (**Fig S2E**). Chromatin accessibility differed among the four groups, from the less accessible constitutive heterochromatic states H1–H6 to the most accessible intergenic states I1–I3, and states E1, E2, and F6 were present in the 5′ and 3′UTRs of genes (**Fig 2E**). Thus, chromatin accessibility, reflecting the potential for transcriptional activity, was broadly associated with chromatin states.

Overall, the chromatin states recapitulated the preferential associations between H2A variants and histone modifications observed in mononucleosomes (**Fig 2A**). Constitutive heterochromatin was enriched in H3K9me1/2, H3K27me1, and H4K20me1, but only H3K9me1 and H2A.W.7 truly represented all six heterochromatic states. The histone modifications H2AK121ub and H3K27me3, which are deposited by PRC1 and PRC2, respectively, were associated with chromatin states F1–F6, and H2A.Z was present in all facultative heterochromatin states. H3K27me3 covered chromatin states F1–F3 and was also observed in H5, while H2AK121Ub marked several euchromatin states. Similarly, H2Bub and H3K36me3 was associated with H2A and H2A.X in all euchromatic states (E1–E11), but all other marks that are usually associated with euchromatin were also distributed over a larger spectrum of states. Contrasting with the strong association between H2A variants and the three major domains of chromatin, much less prominent associations were observed between histone modifications and H3 variants or H2B variants; this finding is in agreement with the results of our biochemical analysis of mononucleosome composition and previous studies (Osakabe et al. 2018; Jiang et al. 2020).

To examine the importance of H2A variants in the definition of chromatin states, we compared the 26 chromatin states identified in our study with the 9 states identified in a previous study that did not include the comprehensive set of histone variants present in Arabidopsis chromatin (Sequeira-Mendes et al. 2014). The blocks of heterochromatic states H1–H6 corresponded to the previously identified states 8 and 9, which were also defined as heterochromatin in (Sequeira-Mendes et al. 2014). States F1–F6 tended to map to the previously defined facultative heterochromatin states 5 and 6, although state F6 was split among three states: 2, 4, and 6. Similarly, although there was a broad correspondence between euchromatin states E1– E11 with states 1, 3, and 7, there were noticeable differences. Overall, several newly defined states were not associated with the corresponding types of chromatin identified in previous studies. In particular, states 2, 4, and 6 contained elements belonging to multiple newly defined chromatin states (**Fig S2B**). Altogether, the 26-states model provides a more refined and coherent classification of elements of chromatin than states defined primarily based on chromatin modifications, pointing to the importance of histone variants in the definition of chromatin states.

To test this idea directly, we calculated chromatin states after excluding either all histone modifications or all histone variants or single families of histone variants. A comparison between the resulting matrices of chromatin states (measured by the Jaccard index) showed that the loss of histone modifications or histone variants affected the definition of the 26 chromatin states to comparable degrees (**Fig 2G**). Removing H2B variants did not affect most chromatin states, except for H1 and I2, which showed high emissions probabilities for H2B variants (**Fig 2A**). By contrast, H3 variants contributed more effectively towards defining specific states, and removing all H2A variants (no H2A) caused the loss of several states marked by a Jaccard index < 0.7 (**Fig 2G**). In summary, histone modifications and histone variants are required to a comparable degree to define biological chromatin states. Among histone variants, H2A variants have the strongest effect on the association with histone modifications to define chromatin states.

### Transposons share distinct CLs that are distinguished by the identity of transposons and their modes of silencing

Constitutive heterochromatin comprised six states that were all enriched to varying degrees in H2A.W.6 and H2A.W.7. These states were further distinguished by their relative enrichment in four histone modifications and H3.1/H3.3 (**Fig 2A**), distinct levels of CHG and CHH DNA methylation (**Fig S3 D,E**), nucleosome density (**Fig S2E**), length (**Fig S2H**), and position relative to the peri-centromere and chromosome arms (**Fig 3A, S3A**, **Table 1**). The relative proximity of heterochromatic states H1–H4 versus H5 and H6 was also revealed by analyzing the closest neighbor relationships between chromatin states (**Fig 2H**). States H2–H4 (occupying pericentromeres) were enriched in transposon genes (TEGs), whereas states H5 and H6 occupied chromosome arms and were primarily associated with transposable element (TE) fragments (**Fig 2F,** and **3A**). TE fragments were methylated by RNA-directed DNA methylation (RdDM), a process involving RNA Pol V (Matzke et al. 2009). Accordingly, we observed enrichment in Pol V ChIP peaks over chromatin states H5 and H6 (**Fig 3B**). These chromatin states were covered by H2A.W.7 (**Fig 2A**) and were typically present in chromosome arms (**Fig 3A**), which are targeted by RdDM (Lorković et al. 2017a; Matzke et al. 2009).

**Fig. 3:**
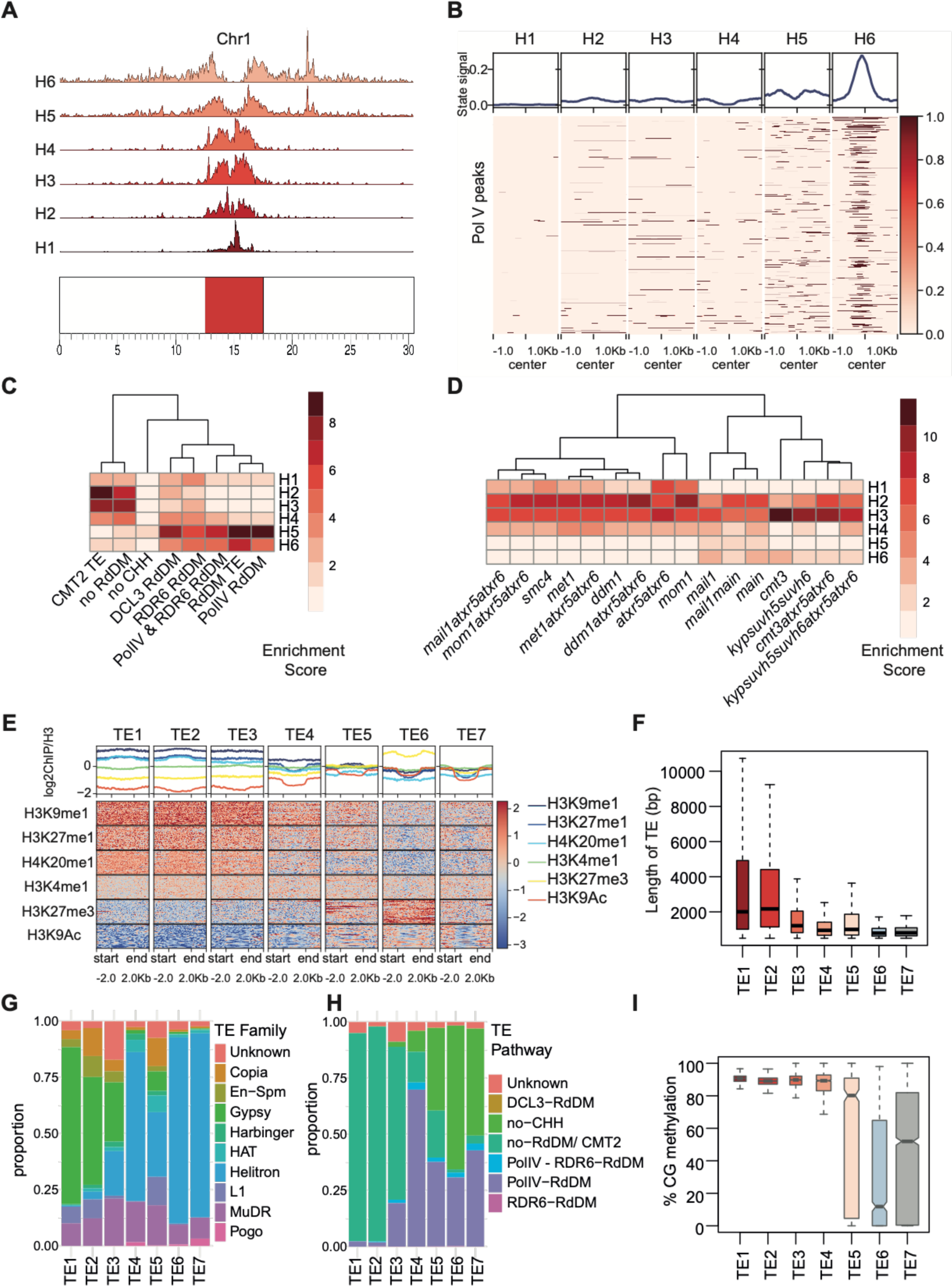
H2A.W.6 and H2A.W.7 variants together describe heterochromatin and repeats. Heterochromatic states can be divided into H2AW6/H2AW7 states (H1–H4) and H2A.W.7- only states (H5–H6). **(A)** Genomic distribution of heterochromatic states (H1–H6) [bottom to top] along Chr1 in *Arabidopsis thaliana*. **(B)** Aggregate heatmap showing the signal of heterochromatic states plotted over Pol V ChIP-seq peaks. **(C)** Heatmap showing the overlap enrichment scores across TEs regulated by different DNA methylation pathways and their association with heterochromatic states. **(D)** Heatmap showing overlap enrichment scores across TE mis-regulated in different heterochromatic mutants and their association with heterochromatic states. **(E)** Aggregate ChIP-seq profiles and heatmaps for various histone modifications across the TE groups. **(F)** Boxplot showing the lengths of TEs in each TE group. **(G, H)** Stacked bar plot showing the proportion of each TE superfamily and TE pathway respectively in each TE group. **(I)** Boxplot showing the CG methylation levels in each TE group based on chromatin states.

**Table 1.**
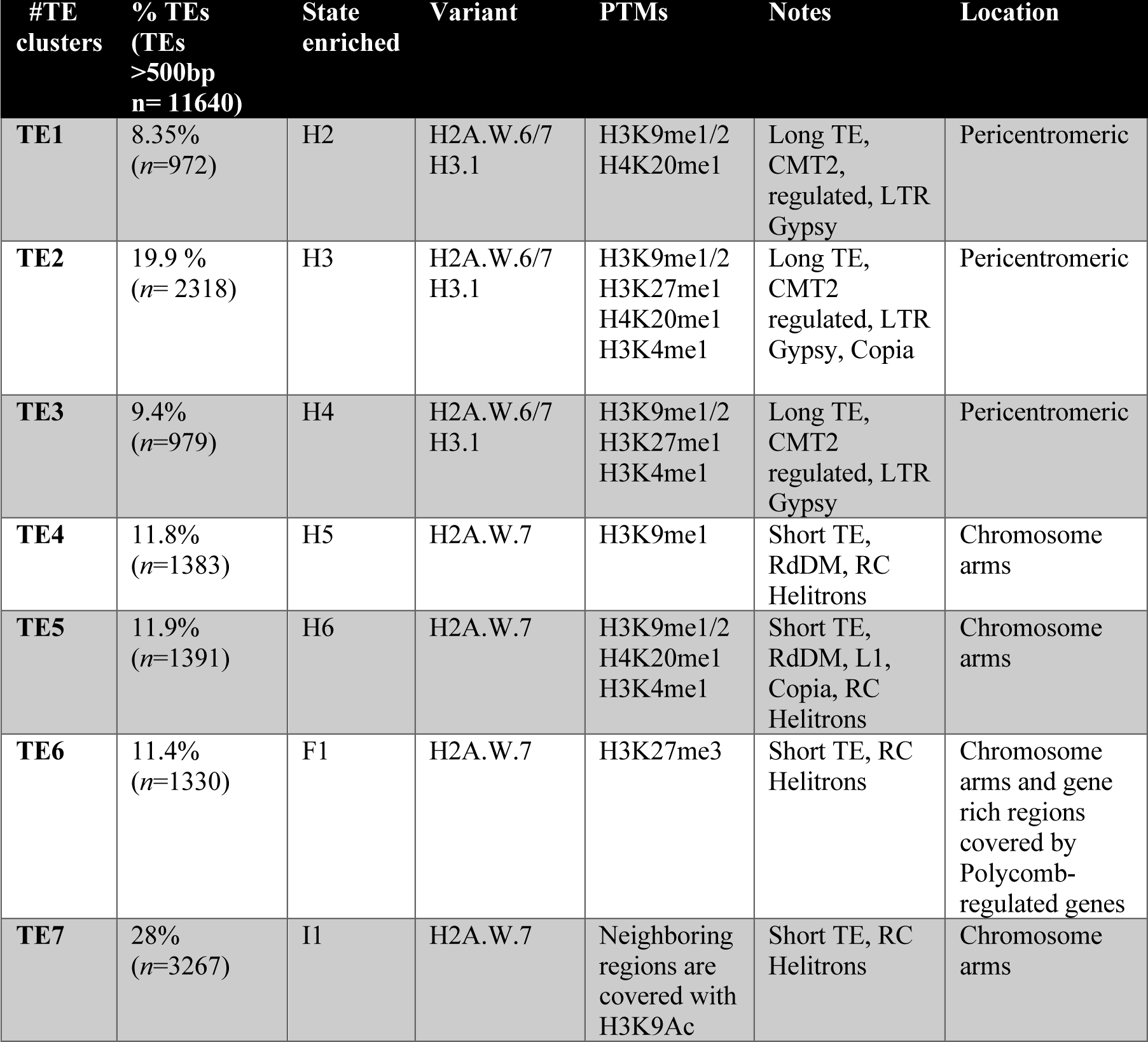
Transposable Elements clusters and their chromatin properties.

We extended our analysis to determine whether there were preferential associations between heterochromatin states H1–H6 and the various mechanisms that deposit DNA methylation (**Fig 3C**) or silence transposons (**Fig 3D**). This confirmed the strong association between RdDM and chromatin states H5 and H6. By contrast, H2 and H3 were not methylated by RdDM but were instead methylated by CHROMOMETHYLASES 2 AND 3 (CMT2 and CMT3), which deposit CHH and CHG methylation (**Fig 3C** and **S3D** and **S3E**). We also observed clear preferential associations between transposon silencing mechanisms and chromatin states (**Fig 3D**), suggesting that chromatin states provide cues that guide silencing mechanisms to their targets.

To establish whether chromatin states associate with each other in a preferential manner, we performed unsupervised hierarchical clustering analysis of the associations between the chromatin states across all transposons. We included only TEs >500 bp long to eliminate fragments of TEs with low mapping ability. We identified seven clusters of TEs, which we named TE1 to TE7 (**Fig S3B**). In each cluster, TEs were primarily enriched in a single chromatin state. The states of constitutive heterochromatin were organized into five clusters (TE1–TE5) over transposons, except for state H1, which did not cover TEs but instead primarily covered repeats at the centers of pericentromeres and were thus likely close to the centromere (**Fig 2F, S2H, S3A, S3B** and **Table 1**). Transposons with no CG methylation were associated with state F1 and were classified as Polycomb group transposons TE6 (**Fig S3B**). Lastly, group TE7 comprised transposons in regions covered with intergenic state I1 (**Fig S3B**).

We also observed marked differences between the seven clusters of TEs in terms of their association with chromatin modifications and histone variants (**Fig 3E, S3F, S3G**), specific TE families (**Fig 3G**), relative proportions amongst all TEs (**Fig S3C**), lengths (**Fig 3F**), and DNA methylation patterns (**Fig 3I** and **S3D–E**). The near exclusive association between TEs from clusters 1–5 and the constitutive heterochromatin states marked by H3K9 methylation confirmed the specific associations among TEs in each cluster, DNA methylation pathways, and enrichment in specific families of TEs. Clusters 1–3, associated with LTR Copia and Gypsy transposons in pericentromeric heterochromatin, were primarily covered by H3.1, H2A.W.6, and H2A.W.7, while TEs from clusters 4 and 5 were located in the arms of chromosomes, covered by H2A.W.7, and methylated by the RdDM pathway. Cluster 6 and 7 primarily contained short Helitrons covered by H2A.W.7 and histone modifications present on protein-coding genes. TE6 transposons were covered with H3K27me3 but no CG methylation, while CG methylation was associated with H3K9Ac on TE7 transposons. TEs from these two clusters were the shortest and located in proximity of protein coding genes (**Fig 2H**).

In conclusion, there were striking associations between TE families and each chromatin state. Such an association was described for the guidance of RdDM by non-coding RNAs on TEs from clusters 4 and 5 (Matzke et al. 2009). Our results generalize this idea and suggest that transposons contain information that guides the deposition of the chromatin marks responsible for their silencing. In addition, we identified a new, large set of associations between TEs and facultative heterochromatin marked by H3K27me3 or intergenic states in euchromatin.

### H2A.Z marks repressed genes in facultative heterochromatin

We applied unsupervised clustering of the relative enrichment of chromatin states to all protein-coding genes of Arabidopsis and observed ten distinct clusters. Unlike TEs, protein-coding genes were occupied by an assembly of chromatin states (**Fig S4A, S4C, FigS5A**). Clustering chromatin states over protein-coding genes revealed that the facultative heterochromatin states F1–F6 formed only three types of associations that represented stereotypical chromatin landscapes (CLs) (**Fig S4A**). Each CL involved the association of one or two chromatin states in the gene body (e.g. F2 and F3 with CL1) with another identical chromatin state upstream and downstream of the gene body (e.g. F1 with Cl1) (**Fig 4A, Fig S4 A, C**). These associations were also supported by neighborhood analysis of chromatin states (**Fig 2H**). Each CL was characterized by a specific combination of chromatin modifications (**Fig 4B**), but CL1–CL3 shared enrichment of H2A.Z on the gene bodies, which represents the hallmark of facultative heterochromatin (**Fig 4B**). The histone modification H2AK121Ub deposited by PRC1 was present in CL1–CL3, although to varying degrees (**Fig 4B**). In CL1, H2AK121Ub associated with H3K27me3 deposited by PRC2 (**Fig 4B**). CL1 comprised ∼70% genes in facultative heterochromatin (**Fig 4A**). These genes shared the lowest expression in the wild type (**Fig 4E**), and many of these genes were expressed in mutants deprived of PRC1 or PRC2 activity (**Fig S4B, Table 2**). In conclusion, genes from CL1 are the genes typically regulated by Polycomb group proteins.

**Fig. 4:**
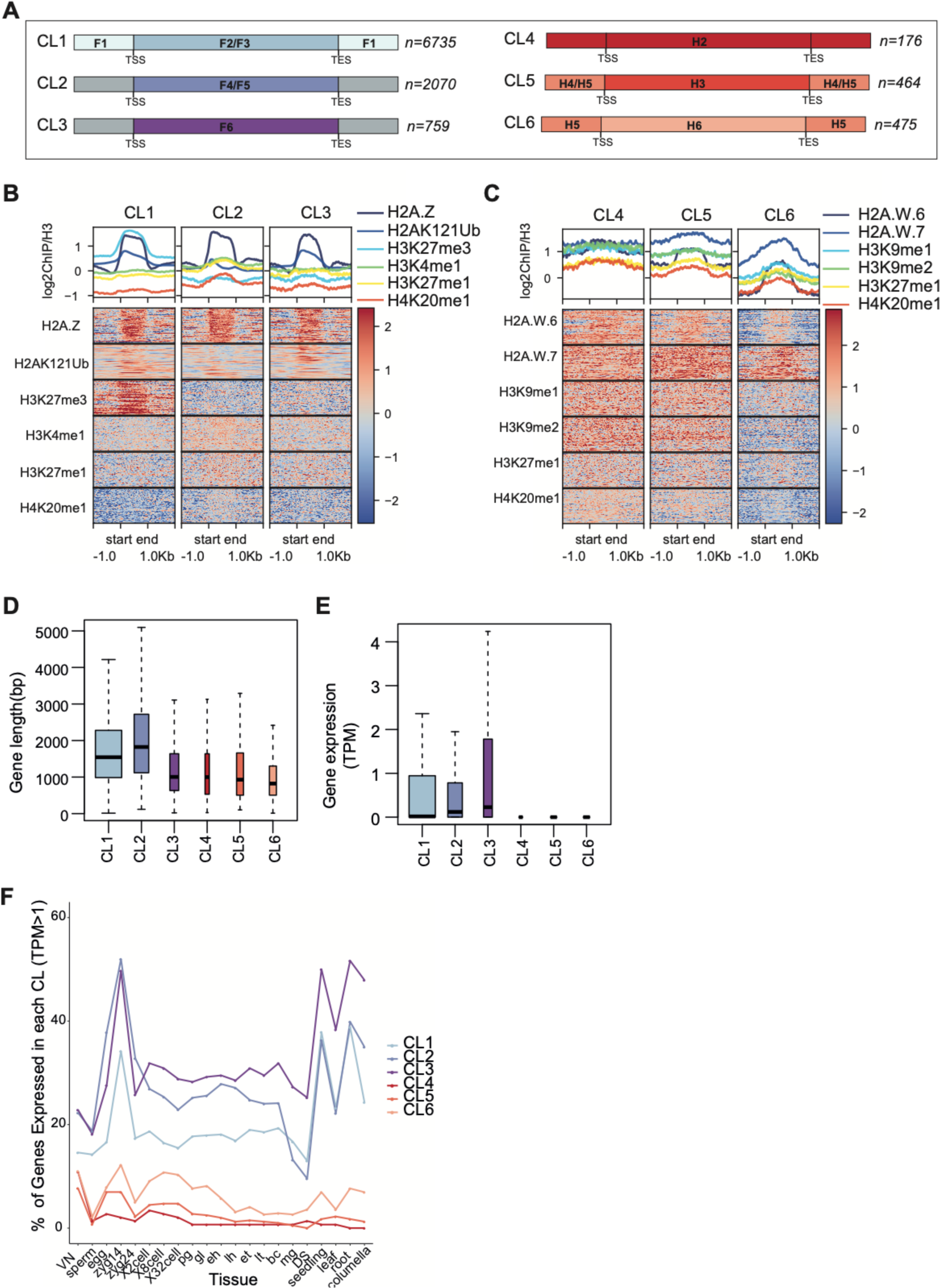
Most H2A.Z in the genome is associated with developmentally repressed genes. There are two types of facultative chromatin states: H2AZ with H3K27me3 (F1–F3) and H2AZ with other repressive marks (F4–F6). **(A)** Schematic representation of six types of repressed genes the chromatin landscapes (CL). [CL1–CL3: Facultative repressed genes and CL4–CL6: Constitutively repressed genes. **(B, C)** Aggregate profile plots and heatmaps showing ChIP-seq signals for histone marks and variants relevant to each CL. **(D)** Boxplot comparing gene lengths across repressed CLs. **(E)** Boxplot comparing gene expression in TPM (transcripts per million) across repressed CLs. **(F)** Line plot showing % Genes expressed (TPM>1) within each CL in different tissue types across various developmental stages in Arabidopsis.

**Table 2.** Specific chromatin landscape of each gene cluster. The proportion of genes, the main histone variant and modifications (PTMs), and specific traits are indicated.

| #Chromatin Landscape | % Protein coding genes | Variant | PTMs | Notes | Gene Ontology terms |
| --- | --- | --- | --- | --- | --- |
| **CL1** | 24.5%<br>(n=6735) | H2A.Z<br>H3.1 | H3K27me3,<br>H2AK121Ub | Strongly repressed by Polycomb, inserted in facultative heterochromatin. Encode developmental genes, miRNAs, lncRNAs | Development, process, and regulation and response related genes |
| **CL2** | 7.5%<br>(n=2070) | H2A.Z<br>H3.1 covering gene body<br>H3.3 only peaks at 3' | H4K20me1,<br>H3K27me1 | Repressed genes independent from Polycomb, inserted in euchromatin. Also include genes encoding snRNAs and snoRNAs | Cell cycle, protein modification related genes |
| **CL3** | 2.7%<br>(n=759) | H2A.Z<br>H3.3 | H2AK121Ub | Short genes weakly repressed by Polycomb and inserted in euchromatin | Defense and response genes |
| **CL4** | 0.7%<br>(n=176) | H2A.W<br>H3.1 | H3K9me1/2 | Genes in pericentric heterochromatin, which are expressed during reproduction. | Regulation of fertilization |
| **CL5** | 1.7%<br>(n=464) | H2A.W<br>H3.1 | H3K9me1/2 | Genes in pericentric heterochromatin | Fertilization |
| **CL6** | 1.8%<br>(n=475) | H2A.W.7<br>H3.1<br>H3.3 | H3K27me1<br>H4K20me1 | Genes in chromosome arms | Pollination |
| **CL7** | 37.0%<br>(n=10140) | H2A/X<br>H2A.Z in 5' | H2Bub<br>H3K36me3<br>H3K4me1<br>H3K4me3 | Highly expressed ubiquitous housekeeping genes, high levels of gene body methylation. Longest genes | Metabolism, housekeeping genes |
| **CL8** | 15.0%<br>(n=4129) | H2A/X<br>H2A.Z in 5' | H3K4me3<br>H3K36me3 | Extended enrichment of H2A.Z inside the gene body and low DNA methylation. Shorter and expressed at lower levels than CL1 genes | Transport, signaling, and response |
| **CL9** | 3.9%<br>(n=1057) | H2A.Z | H2AK121Ub | Short genes devoid of intron and gene body methylation. Low levels of RNA Pol II. Resemble genes in clusters CL5 and CL7 | Response to karrikin<br>Only one GO terms found for this group |
| **CL10** | 5.2%<br>(n=1440) | H2A/X | H3K4me1 high<br>H3K4me3 low<br>H3K36me3 low<br>H4K20me1<br>H3K27me1 | Lowest levels of RNA Pol II, transcripts, and chromatin accessibility and high levels of gene body methylation. Similar to CL6 genes. | Cell cycle, chromosome segregation |

CL2 and CL3 comprised genes covered by H2A.Z marked by H2AK121Ub and H3.3 (Fig. 4B, S4E and Table 2). Whereas genes from CL1 were bordered by regions enriched in H3K27me3, genes within CL2 also carried marks present in constitutive heterochromatin and were occupied and surrounded by regions enriched in H3K4me1 and H3K27me1 (**Fig 4B**). Genes within CL3 were shorter than those in CL1 and CL2 (**Fig 4D**) and were enriched in H3.3 compared to other genes in constitutive heterochromatin (**Fig S4E**).

In addition to the 9564 repressed genes covered by CL1–CL3 typical of facultative heterochromatin (dominated by the presence of H2A.Z), 1115 protein-coding genes were not expressed and were covered primarily by heterochromatic states (**Fig 4A, C, E,** and **S4C–D, Table 2**). These genes were covered by CL4–CL6, which were distinguished by constitutive heterochromatin states present at their gene bodies (**Fig 4A** and **S4C**) and were preferentially enriched for H2A.W.7 versus H2A.W6 and H3.1 versus H3.3 in their gene bodies (**Fig S4D– E**). These genes were preferentially associated with pericentric heterochromatin (CL4 and CL5) and were surrounded by H2A.W.7 and H3K9me (**Fig 4C**) or with heterochromatin islands in chromosome arms (CL6).

We conclude that constitutive heterochromatin hosts genes with three types of chromatin landscapes dominated by H3.1 in association with H2A.W and H3K9me. By contrast, H2AK121Ub is the main marker of facultative heterochromatin, which comprises three types of chromatin landscapes across genes controlled by Polycomb repressive complexes.

### Active genes share one of four stereotypical chromatin landscapes

All 16,776 expressed protein-coding genes shared some of the euchromatin states (E1–E11), which assembled into four chromatin landscapes (CL7–CL10; **Fig S5A, Table 2**). These associations were also supported by neighborhood analysis of chromatin states (**Fig 2H**). The E1 state enriched in H2A.Z (**Fig 2A**), the only state common to all euchromatin genes, marked the 5′UTR, except CL9, where E1 also marked the entire gene body (**Fig 5A and S5A**). CL9 was most similar to CL1–CL3 and occupied genes with gene bodies covered by H2AK121Ub, but it was associated with euchromatin marks H3K4me3, H3K36me3, and H3 acetylation. All other CLs comprised an assembly of euchromatic states arranged in the 5′ to 3′ direction from E1–E9 for cluster 1, E1–E3, E8 for cluster 2, and E1 followed by E10/E11 for CL10 (**Fig 5A** and **S5A**). All euchromatic CLs were characterized by gene bodies surrounded by the intergenic state I1 (except for I3, which was located upstream of the 5′ end for CL8) enriched in H3K9Ac and H3K14Ac and with the highest chromatin accessibility (**Fig 2A, 2E**). In genes covered by CL7–CL10, the transcription end site (TES) was also routinely followed by a 3′UTR marked by the state I2 characterized by enrichment in histone variant H2B.1 and H2B.2 (**Fig 2A, 2F, 5A, S5A, S6B**).

**Fig. 5:**
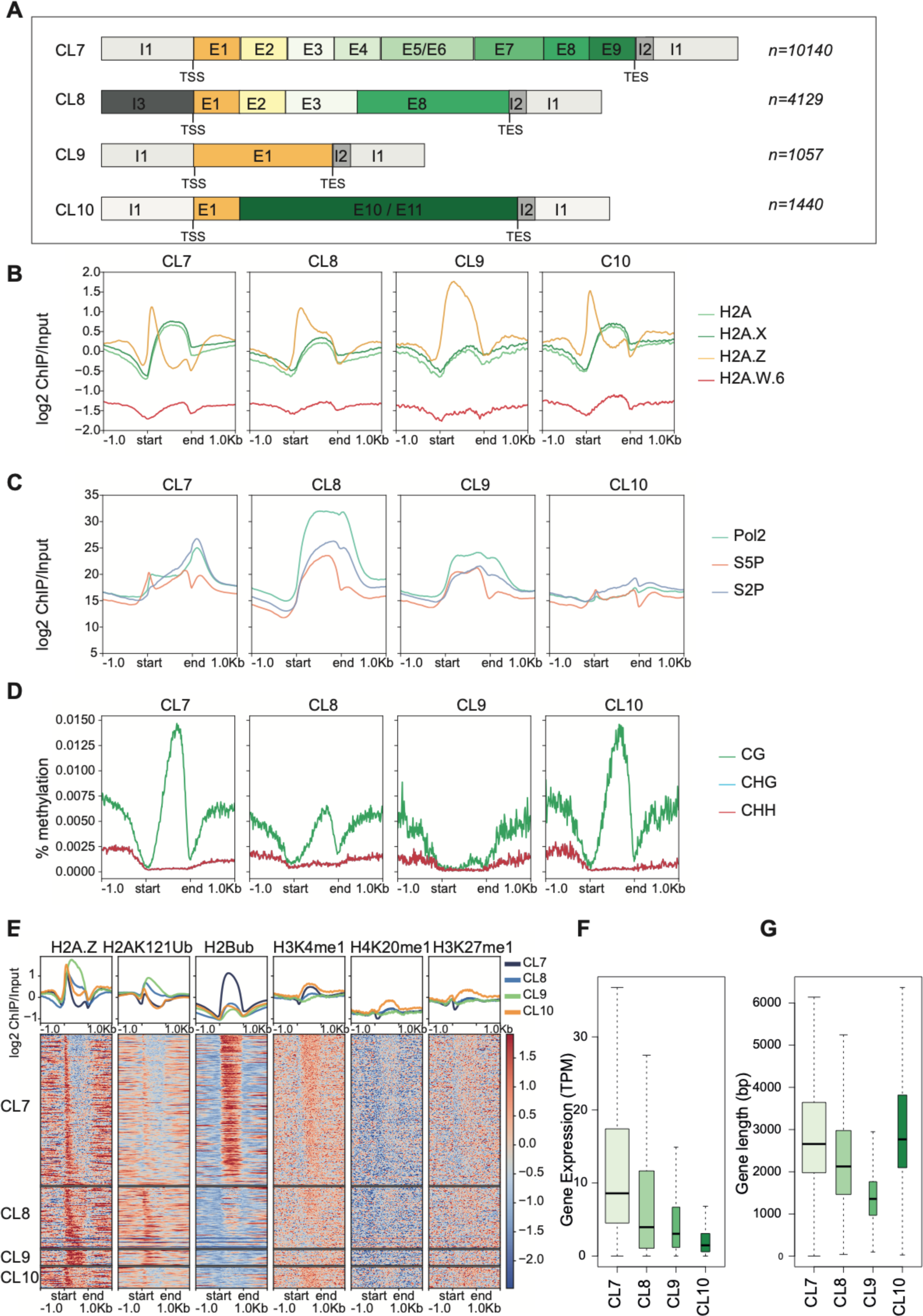
Transcribed chromatin states assemble in a stereotypical fashion across active genes. Euchromatic states assemble in a stereotypic fashion across expressed protein-coding genes. **(A)** Schematic representation of states occupying the four types of euchromatic gene clusters CL7–CL10 [CL7: Regular active genes; CL8: Low CG methylated genes; CL9: Short H2AZ genes; and CL10: Bivalent genes]. **(B–D)** Aggregation profile plots across the four types of active gene CLs of histone H2A variants (B), Pol II occupancy (C), and DNA methylation levels (D). **(E)** Aggregation profile and heatmap showing log2(ChIP/Input) for chromatin marks that differentiate across the four gene CLs. **(F, G)** Boxplots showing gene expression (F) and gene length (G) across active gene clusters.

Genes from all euchromatic CLs shared similar profiles of H3K4me3 just after the TSS overlaid with a broad asymmetrical enrichment of H3K36me3 declining towards the 3′ end (**Fig S5B**). The H3K4me3 profile overlaid a similar peak of H2A.Z, H3K9Ac, and H3K14Ac at the 5′ end, which mark the presence of the preinitiation complex (Leng et al., 2020). One exception was genes with CL9, which showed a symmetrical enrichment in these marks parallel to their symmetrical organization of chromatin. Overall, each euchromatic CL exhibited distinctive features relative to the genomic profiles of H2A.Z, H2A.X, and replicative H2A (**Fig 5B**). Genes with CL7 and CL10 showed a strong peak of H2A.Z at the 5′ end, while this peak was extended towards the 3′ end in genes with CL8 and CL9. In opposition to the enrichment of H2A.Z, CL7 and CL10 shared the strong presence of CG methylation towards the 3′ of the gene body, while this enrichment was reduced in CL8 and CL9 (**Fig 5C**). In addition, we observed a gradual enrichment of H3.3 towards the 3′ ends of all euchromatic genes, as reported previously (Wollmann et al. 2017). In addition, in general, the variant H2B.3 was located over gene bodies, while H2B.1 and H2B.2 marked the beginning of the 3′UTRs (**Fig S6B**).

The characteristics of CL8–CL10 are detailed in Table 2. These CLs lacked the states E5, E6, and E7 and were present in genes with lower expression levels, and except for CL10, in genes that were shorter than those covered with CL7 (**Fig 5A, 5F,** and **5G**). Two-thirds of euchromatin genes were covered with CL7 and were (on average) the longest and most strongly transcribed (**Fig 5A, 5F,** and **5G**). These genes covered with CL7 were marked by strong enrichment of H2Bub from the 5′ to 3′ end, which was not observed in other euchromatic genes (**Fig 5E**). The transition to the gene bodies was marked by the replacement of H2A.Z by H2A and H2A.X associated with H2Bub and H3K36me3. The loss of the latter histone modifications marked the transition to states E8 and E9, with a return of H3K9Ac that marked the terminators.

In conclusion, all CLs of euchromatin showed an asymmetrical modular organization that primarily reflected the distribution of H2A and H2A.X in association with H3.3, H2B.3, and H3K36me3. This organization contrasts with the symmetric arrangement of chromatin over CLs associated with gene repression.

### Chromatin landscapes determine a range of transcriptional activity

CL7 and CL10 showed a peak of S5P Pol II at the 5′ ends of genes and a peak of S2P towards the ends of genes, although these features were much less pronounced in CL10 (**Fig 5C**). By contrast, these peaks were absent in CL8 and CL9, with a much higher and uniform profile of enrichment of Pol II covering the entire transcriptional unit from the 5′ to 3′ end (**Fig 5C**). Surprisingly, the coupling of CL7 and CL10 in opposition to CL8 and CL9 was not reflected in the range of gene expression, which was strongest in CL7 genes and diminished from CL8 to CL10 genes (**Fig 5F**). Similarly, compared to the high levels of expression related to CL7 and the lowest levels of expression related to CL10, the transcript levels were correlated with the enrichment of H2A.X, H2A, and several modifications over the gene bodies (**Fig 5F** and **S6**).

To identify features that were directly correlated with the level of transcription, we compared the enrichment of each mark across gene quartiles based on their transcript levels. In each CL, we observed a clear positive correlation between transcript levels, chromatin accessibility revealed by ATAC-seq (**Fig S7**) and the levels of enrichment in RNA Pol II and its phosphorylated forms S5P (paused) and S2P (elongating) (**Fig S8**). Each CL was associated with a distinct profile of RNA Pol II and its phosphorylated forms, with patterns of peaks and troughs unique for each CL. The overall shape of the pattern did not vary between quartiles which correlated positively with increased enrichment of RNA Pol II. Remarkably, we did not observe such a marked correlation between transcript levels and the enrichment of H2A and H3 variants or histone H3 and H4 modifications or gene body methylation, except for weak correlations for CL8 (**Fig S7, S9–S10**). The exception was H2Bub enrichment, which was positively associated with the levels of transcripts in each CL (**Fig S8–S10**). In conclusion, within the set of genes associated with a particular CL, the chromatin composition did not vary with transcript levels. Moreover, each CL was associated with a specific range of transcript levels and a distinct profile of RNA Pol II, suggesting that euchromatic CLs are set up in a stereotypical manner that constrains RNA Pol II activity to a range of levels.

## Discussion

### Histone variants drive the overall organization of chromatin into CLs

We demonstrated that histone modifications and histone variants subdivide the intergenic space, the non-protein-coding space, and the protein-coding genic space of the genome into 26 chromatin states. This refines the previous definition of nine chromatin states in Arabidopsis (Sequeira-Mendes et al. 2014). Our analysis indicated that histone variants are as impactful as histone modifications in defining chromatin states. *In silico* analyses showed that H2A variants are most important in defining the broad groups of chromatin states. These chromatin states include constitutive heterochromatin defined by H2A.W and H3K9me1; facultative heterochromatin defined by H2A.Z carrying K121Ub, with a strong association with H3K27me3; and euchromatin, primarily occupied by H2A and H2A.X in conjunction with H3K4me3, H3K36me3, and histone acetylation. These strong associations between H2A variants and histone modifications are supported by the results of biochemical analysis (our analysis and (Osakabe et al. 2018).

The mechanisms that drive the stereotypical associations between H2A variants and specific modifications remain unclear in the case of H2A.W, H2A.X, and replicative H2A, but we have some clues about the role of H2A.Z and the Polycomb writers PRC1 and PRC2. H2A.Z is the major carrier of H2AK121Ub deposited by PRC1, as indicated by their co-occurrence with chromatin states as well as previous biochemical analyses (Osakabe et al. 2018). *In vitro* biochemical assays also showed that PRC2 strongly prefers arrays of H2A.Z as a substrate(Y. Wang et al. 2018), thus supporting the link between H2A.Z and H3K27me3 (Carter et al. 2018). Overall, our observations suggest that chromatin writers and erasers likely operate with different affinities and efficacies based on the histone variants in the nucleosome. This is also illustrated by the anticorrelation between H2Bub and H2A.Z resulting in the inhibition of the ubiquitin ligase by the H2A.Z tail (Surface et al. 2016) and by the link between H3K27 monomethylation of its preferred substrate H3.1(Jacob et al. 2014). We also observed a relationship between H3.3 and H3K36me1/2, pointing to a possible preference of H3.3 for the deposition of H3K36me or for the demethylation of H3K36me3. Although a general enrichment of H3.1 in constitutive heterochromatin was also noted previously, H3.1 and H3.3 do not differentiate the three main compartments of chromatin but tend to differentiate gene bodies among the different CLs. H2B variants define specific states: H1 close to the centromere, and I2 marking the 3′UTRs of expressed genes. However, their role in defining these states is minor compared to roles of specific H2A variants, whose presence or absence provides global discrimination of the three major domains of the genome and their subdivisions.

### Chromatin states are arranged into stereotypical CLs on transposons and genes

A major advance using our approach resulted from clustering chromatin states over transposons and genes. As summarized in Tables 1 and 2, the Arabidopsis genome is occupied by only a limited number of distinct CLs. In addition to these we expect two additional CLs including the centromere, which was recently described (Naish et al. 2021), and the ribosomal DNA arrays that were not studied here due to lack of unique mapping ability of these regions (Sims et al. 2021). We described seven CLs that occupy transposons, while protein-coding genes are occupied by ten contrasting CLs. The symmetrical organization of chromatin states over CLs invariably marks low or no expression, while modular directional assemblies of chromatin states over CLs are associated with expressed genes.

Of the 11640 annotated TEs, 7043 are covered by CLs in constitutive heterochromatin. Among these, most TEGs are present in CLs in the pericentromeres and are dominated by H2A.W and H3K9me, which act in synergy to silence transposons in the pericentromeric regions (Bourguet et al. n.d.) Broadly, we observed strong associations between the identity of transposons and specific silencing mechanisms. This provides concrete evidence for the selective adaptation of transposons to distinct silencing machineries (Bourguet et al. n.d.). Our analysis supports and extends the idea that different mechanisms act in distinct territories of heterochromatin. Our results suggest that TEs segregate in distinct territories that recruit different actors of silencing. We also determined that 4597 TEs (clusters 2 and 3), mostly transposon fragments and short TEs, are present in intergenic regions. Hence, a large fraction of annotated TEs that are associated with CLs typically associate with protein-coding genes, suggesting that TEs play a pervasive role in transcriptional regulation; this represents one of only a few examples of this concept reported to date (Quadrana 2020).

In Arabidopsis seedlings, circa 40% of protein-coding genes are repressed and covered by three types of CLs typical of facultative heterochromatin; 70% of these genes are present in CLs dominated by domains targeted by both PRC1 and PRC2 and represent Polycomb complex-controlled genes. The remaining 30% of genes are isolated and covered only by the marks typical of constitutive heterochromatin in addition to H2AK121Ub (CL2 and CL3) or are present in constitutive heterochromatin (CL4 to CL6). The other protein-coding genes are expressed to varying degrees and are covered with four distinct CLs in which the first few nucleosomes are enriched in H2A.Z and H3K4me3. CL9 occupies short genes covered by H2A.Z that are expressed at low levels, while most genes with CL7 or CL8 are largely occupied by replicative H2A and H2A.X. The central module of CL7 hosts gene body methylation and is surrounded by a module of four states that structures the 5′ end, while a module of two states marks the 3′ ends of genes. CL8 is similar to CL7 but lacks the central module and thus shows very low gene body methylation. By contrast, approximately 1000 genes with the lowest expression levels are occupied by CL10, with a specific core state occupied by high gene body methylation and constitutive heterochromatin marks H3K27me1 and H4K20me1.

We conclude that CLs define distinct environments associated with transcriptional silencing, repression, or activity. These CLs are differentiated by different patterns of enrichment of H2A variants, H3 variants, and histone modifications. The limited number of the CLs indicates that they are produced by via the orchestration of dedicated mechanisms, which depend on the coordinated activity of chromatin writers/erasers as well as proteins that regulate the deposition and eviction of histones. While in plants chaperones specific for H3 deposition are well studied, chaperones for each type of H2A variant have not been identified (Nie et al. 2014), several remodelers are involved in the exchange of H3.1 with H3.3 (H. Wang et al. 2018), H2A.Z with replicative H2A (Aslam et al. 2019), the eviction of H2A.Z (Aslam et al. 2019), or the deposition of H2A.W (Osakabe et al. 2021).We propose that these remodelers participate in determining the architecture of the different CLs together with the chaperones that control the specific deposition of H3.1 or H3.3.

### CLs determine the range of transcriptional activities of protein-coding genes

The well-characterized genome-wide association between transcript levels and the enrichment of several histone modifications and histone variants suggests that epigenetic modifications and/or the deposition of histone variants either modify transcription or are a consequence of transcription(Leng et al. 2020). Histone H3 and H4 acetylation neutralize the charge of lysine residues, destabilize nucleosomes, and facilitate the promotion of transcription(Leng et al. 2020), but the role of H3K4me3 and H3K36me3 in promoting transcription is less clear (Talbert and Henikoff 2021). For each CL associated with expressed genes, we discovered the surprising lack of correlation between transcript levels and the levels of histone variants, gene body DNA methylation, and H3 and H4 modifications. The range of transcription in each CL is not narrow and declines from CL7 to CL10, and there is some overlap between the ranges of these four CLs. We believe that this explains the correlations observed between transcript levels and the enrichment of chromatin marks when the entire pool of protein-coding genes was included. However, it is clear that each CL is associated with a distinct profile of RNA Pol II. We thus conclude that each CL (comprising a fixed pattern of enrichment of chromatin modifications and histone variants) enables a specific regime of RNA Pol II activity, resulting in a specific range of transcripts produced. H2bUb is the only histone modification studied here that was consistently affected in all CLs by transcript levels. This could be explained by the fact that H2Bub is a hub for the recruitment of factors involved in transcriptional elongation through interactions with H3K4me3 (Leng et al. 2020) and associations with H3K36me3 writers (McGinty et al. 2008; Zhao et al. 2019). The association of HUB1 with transcription elongation factors highlights the importance of this modification for productive transcriptional elongation (Antosz et al. 2017; Pfab, Breindl, and Grasser 2018). The ubiquitination of H2B is strongly inhibited by the H2A.Z tail and the specific acidic patch of H2A.Z in mammals (Wojcik et al. 2018). The strong opposition of H2A.Z and H2Bub in Arabidopsis supports the likely conservation of the inhibition of ubiquitination of H2B by the H2A.Z tail and the specific acidic patch shown in mammals (Wojcik et al. 2018). H2Bub stimulates the deposition of H3K36me3 and productive transcription elongation and thus appears to function as a relay between the CL and the activity of RNA Pol II. The effect of transcription on H2Bub observed in our study also suggests a feedback loop that could involve H3K4me3 (Leng et al. 2020).

Our findings argue in favor of a deterministic nature of CLs that would inform the transcriptional activity associated with genes or transposons in Arabidopsis. Although the organization of the transcriptional units of plants is somewhat different from that of yeast or mammals, chromatin modifications and histone variants are strongly conserved across a large range of eukaryotes (Grau-Bové and Sebé-Pedrós 2021), suggesting that comparable CLs will be identified in a broad range of organisms. If so, the parsimonious nature of CLs and their capacity to instruct the orientation and level of transcriptional activity imply the existence of a genetic code that shapes the organization of CLs. Like the preferred sequences that direct chromatin remodelers to define the structure of the TSS (Raisner et al. 2005; J. Wang et al. 2019) and the regular spacing of nucleosomes (Li et al. 2014), the sequences of genes that share a common CL share common features that should now be deciphered. Finding the cis-code that guides the complex machinery that builds the CL will be the next step in decoding the relationship between the genetic code and the code provided by the CLs of histone variants and modifications. Perhaps this code could ultimately be used to engineer chromatin organization in synthetic eukaryotic systems.

## Methods

### Generation of antibodies, isolation of nuclei, MNase digestion, immunoprecipitation, SDS-PAGE and Western blotting

Antibodies against H2A.Z.11 (KGLVAAKTMAANKDKC) and H2A.2 (CPKKAGSSKPTEED) peptides were raised in rabbits (Eurogentec) and purified by peptide affinity column. Purified IgG fractions were tested for specificity on nuclear extracts from WT and knock out lines or with overexpressed histone variants.

For MNase digestion followed by immunoprecipitation, nuclei were isolated from 4 grams of 2-3 weeks old leaves and the procedure as described in (Lorković et al. 2017) was followed. Isolated nuclei were washed once in 1 ml of N buffer (15 mM Tris-HCl pH 7.5, 60 mM KCl, 15 mM NaCl, 5 mM MgCl2, 1 mM CaCl2, 250 mM sucrose, 1 mM DTT, 10 mM ß- glycerophosphate) supplemented with protease inhibitors (Roche). After spinning for 5 min at 1,800 × *g* at 4°C nuclei were re-suspended in N buffer to a volume of 1 ml. Twenty microliters of MNase (0.1 u/µl) (SigmaAldrich) were added to each tube and incubated for 15 min at 37°C and during the incubation nuclei were mixed 4 times by inverting the tubes. MNase digestion was stopped on ice by addition of 110 µl of MNase stop solution (100 mM EDTA, 100 mM EGTA). Nuclei were lysed by addition of 110 µl of 5 M NaCl (final concentration of 500 mM NaCl). The suspension was mixed by inverting the tubes and they were then kept on ice for 15 min. Extracts were cleared by centrifugation for 10 min at 20,000 × *g* at 4°C. Supernatants were collected and centrifuged again as above. For each immunoprecipitation extract, an equivalent of 4 g of leaf material was used, usually in a volume of 1 ml. To control MNase digestion efficiency, 100 µl of the extract were kept for DNA extraction. Antibodies, including non-specific IgG from rabbit, were bound to protein A magnetic beads (GE Healthcare) and then incubated with MNase extracts over night at 4°C. Beads were washed two times with N buffer without sucrose, containing 300 mM NaCl, followed by three washes with N buffer containing 500 mM NaCl. Beads were re-suspended in 150 µl of 1 × Laemmli loading buffer in 0.2 × PBS. Proteins were resolved on 15% SDS-PAGE, transferred to a Protran nitrocellulose membrane (GE Healthcare) and analyzed by Western blotting using standard procedures. The blots were developed with an enhanced chemiluminescence kit (Thermo Fischer Scientific) and signals acquired by a ChemiDoc instrument (BioRad). All primary histone variant-specific and H3 marks-specific antibodies were used at 1 µg/ml dilution. H3 specific antibody was used at 1:5,000 dilution. Rat anti-HA antibody (Roche 3F10) was used at 1:2,000 dilution. Secondary antibodies were goat anti-rabbit IgG (BioRad) and goat-anti rat IgG (SigmaAldrich), both at a 1:10,000 dilution.

### Mass spectrometry

For mass spectrometry immunoprecipitated nucleosomes were resolved on 4–20% gradient gels (Serva) and silver-stained. Histone H3 bands were excised, reduced, alkylated, in-gel trypsin and LysC digested, and processed for MS. The nano HPLC system used was a Dionex UltiMate 3000 HPLC RSLC nano system (Thermo Fisher Scientific, Amsterdam, Netherlands) coupled to a Q Exactive mass spectrometer (Thermo Fisher Scientific, Bremen, Germany), equipped with a Proxeon nanospray source (Thermo Fisher Scientific, Odense, Denmark). Peptides were loaded onto a trap column (Thermo Fisher Scientific, Amsterdam, Netherlands, PepMap C18, 5 mm 3 300 mm ID, 5 mm particles, 100 Å pore size) at a flow rate of 25 ml/min using 0.1% TFA as the mobile phase. After 10 min, the trap column was switched in line with the analytical column (Thermo Fisher Scientific, Amsterdam, Netherlands, PepMap C18, 500mm 3 75 mm ID, 2 mm, 100 Å). Peptides were eluted using a flow rate of 230 nl/min and a binary 2-h gradient. The gradient starts with the mobile phases: 98% solution A (water/formic acid, 99.9/0.1, v/v) and 2% solution B (water/acetonitrile/formic acid, 19.92/80/0.08, v/v/v), increases to 35% of solution B over the next 120 min, followed by a gradient in 5 min to 90% of solution B, stays there for 5 min and decreases in 5 min back to the gradient 98% of solution A and 2% of solution B for equilibration at 30°C. The Q Exactive HF mass spectrometer was operated in data-dependent mode, using a full scan (m/z range 380–1500, nominal resolution of 60 000, target value 1E6) followed by MS/MS scans of the 10 most abundant ions. MS/MS spectra were acquired using a normalized collision eneergy of 27%, an isolation width of 1.4 m/z, and a resolution of 30.000, and the target value was set to 1E5. Precursor ions selected for fragmentation (exclude charge state 1, 7, 8, >8) were put on a dynamic exclusion list for 40 s. Additionally, the minimum AGC target was set to 5E3 and intensity threshold was calculated to be 4.8E4. The peptide match feature was set to preferred, and the exclude isotopes feature was enabled. For peptide identification, the RAW files were loaded into Proteome Discoverer (version 2.1.0.81, Thermo Scientific). The resulting MS/MS spectra were searched against A. thaliana histone H3 sequences (7 sequences; 951 residues) using MSAmanda v2.1.5.8733 (Dorfer et al., 2014). The following search parameters were used: Beta-methylthiolation on cysteine was set as a fixed modification, oxidation on methionine, deamidation on asparagine and glutamine, acetylation on lysine, phosphorylation on serine, threonine, and tyrosine, methylation and di-methylation on lysine and threonine, tri-methylation on lysine, and ubiquitinylation on lysine were set as variable modifications. The peptide mass tolerance was set to ±5 ppm, and the fragment mass tolerance was set to ±15 ppm. The maximal number of missed cleavages was set to 2. The result was filtered to 1% FDR at the peptide level using the Percolator algorithm integrated in Thermo Proteome Discoverer. The localization of the phosphorylation sites within the peptides was performed with the tool ptmRS, which is based on the tool phosphoRS (Taus et al., 2011).

Peptides diagnostic for H3.1 and H3.3 covering positions K27 and K36 (see **Fig S1D**) were used for the analysis of modifications. All peptides covering these two positions were selected in H3.1 and H3.3 immunoprecipitation samples and analyzed for the presence of modifications with threshold for modification probability set to 95% or higher. Relative modification levels were expressed as number of modified peptides (**Fig S1F**) divided by total number of peptides (**Fig S1E**) that were measured for each lysine position resulting in total modification levels for H3.1 and H3.3 (see **Fig 1D**). We also analyzed the same data by splitting H3.1 and H3.3 specific peptides in each immunoprecipitation and obtained highly similar trends for H3.1 and H3.3 irrespective of whether they were precipitated with H3.1 or H3.3 (**Fig S1**). Histone acetylation was analyzed by selecting all peptides covering indicated positions and expressed as relative acetylation levels in each immunoprecipitation without differentiating H3.1 and H3.3 variants (**Fig S1H and S1I**). We also analyzed H3K9, H3K14 and H3K18 acetylation from peptides derived from transgenic copy alone because these data reflect modification levels of each H3 variant and obtained highly similar levels (**Fig S1J**) as without differentiation between H3.1 and H3.3.

### Plant material for ChIP-seq

Col-0 Wild type (WT) *Arabidopsis thaliana* seeds were stratified at 4°C in the dark for three days. Seeds were grown on ½ MS sterilized plates in the growth chamber under long day (LD) conditions (21°C; 16 h light/8 h dark). After 10-days seedling tissue was freshly harvested.

### Chromatin immunoprecipitation (ChIP)

For ChIP nuclei isolation was performed using 10-day seedlings from WT Col-0. The procedure for nuclei isolation and chromatin immunoprecipitation was performed as described in (Osakabe et al. 2021). Freshly harvested tissues (0.3 g of tissue was used for each immunoprecipitation) were fixed with 1% formaldehyde for 15 min and the cross-linking reaction was stopped by the addition of 125 mM glycine. Crosslinked tissues were frozen and ground with a mortar and pestle in liquid nitrogen to obtain a fine powder. Ground tissues were resuspended in M1 buffer (10 mM sodium phosphate pH 7.0, 100 mM NaCl, 1 M hexylene glycol, 10 mM 2-mercaptoethanol, and protease inhibitor (Roche)), and the suspension was filtered through miracloth. Nuclei were precipitated by centrifugation and washed six times with M2 buffer (10 mM sodium phosphate pH 7.0, 100 mM NaCl, 1 M hexylene glycol, 10 mM MgCl2, 0.5% Triton X-100, 10 mM 2-mercaptoethanol, and protease inhibitor), and then further washed once with M3 buffer (10 mM sodium phosphate pH 7.0, 100 mM NaCl, 10 mM 2-mercaptoethanol, and protease inhibitor). Nuclei pellets were rinsed and resuspended in a sonication buffer (10 mM Tris-HCl pH 8.0, 1 mM EDTA, 0.1% SDS, and protease inhibitor). Nuclei were sonicated with a Covaris E220 High Performance Focused Ultrasonicator for 15 min at 4°C (Duty factor 5.0; PIP 140.0; Cycles per Burst 200) in 1 ml Covaris milliTUBE. After chromatin shearing, the debris were removed by centrifugation and the solutions containing chromatin fragments were diluted with three times the volume of ChIP dilution buffer (16.7 mM Tris-HCl pH 8.0, 167 mM NaCl, 1.2 mM EDTA, 1.1% Triton X-100, 0.01% SDS, and protease inhibitor). After dilution, protein A/G magnetic beads (50 µl for one gram of tissue; Thermo Fisher Scientific) were added to sheared chromatin and incubated at 4°C for 1 hour with rotation. Pre-cleared samples were collected and incubated with 5 µg of anti-H3, anti-H2A.X.3/5,anti-H2A, anti-H2A.Z.9/11, anti-H2A.W.6, anti H2A.W.7, anti-H1, H3K36me3 (Abcam, ab9050), anti-H3K27me3 (Millipore, 07-449), anti-H3K4me3,anti- H3K4me1, anti-H3K27me1, anti-H4K20me1, anti-H3K9me1 or anti-H3K9me2 (Abcam, ab1220) [J1] antibodies at 4°C overnight with rotation. After incubation, samples were mixed with 30 µl of protein A/G magnetic beads, incubated at 4°C for 3 hours with rotation, washed twice with low salt buffer (20 mM Tris-HCl pH 8.0, 150 mM NaCl, 2 mM EDTA, 1% Triton X-100, and 0.1% SDS), once with high salt buffer (20 mM Tris-HCl pH 8.0, 500 mM NaCl, 2 mM EDTA, 1% Triton X-100, and 0.1% SDS), once with LiCl buffer (10 mM Tris-HCl pH 8.0, 1 mM EDTA, 0.25 M LiCl, 1% IGEPAL^®^ CA-630, and 0.1% deoxycholic acid), and twice with TE buffer (10 mM Tris-HCl pH 8.0 and 1 mM EDTA). Immunoprecipitated DNA was eluted by adding 500 µl of elution buffer (1% SDS and 0.1 M NaHCO3), incubated at 65°C for 15 min, and mixed with 51 µl of reverse crosslink buffer (40 mM Tris-HCl pH 8.0, 0.2 M NaCl, 10 mM EDTA, 0.04 mg/ml proteinase K (Thermo Fisher Scientific)). The reaction mixture was then incubated at 45°C for 3 hours and subsequently at 65°C for 16 hours. After crosslink reversal, DNA was treated with 10 µg of RNaseA (Thermo Fisher Scientific), incubated at room temperature for 30 min, and purified with a MinElute PCR purification kit (Qiagen).

### ChIP-seq library prep and data analysis

For ChIP- seq, libraries were prepared with an Nugen Ovation Ultralow V2 DNA-Seq library prep kit (NuGen)following the manufacturer’s instructions. The libraries were sequenced with an Illumina Hiseq 2000 to generate single-end 50 bp reads. For alignment and quality check of sequenced samples, bedtools v2.27.1 was used to convert the raw BAM files to fastq. FastQC v0.11.8(Andrews 2010) (http://www.bioinformatics.babraham.ac.uk/projects/fastqc/) was used to generate quality reports for all sequencing data. Reads were filtered and then aligned to the TAIR10 *Arabidopsis* genome using Bowtie2 (Langmead and Steven L Salzberg 2013) with default settings. Deeptools v3.1.2(Ramírez et al. 2016) was used to examine correlations between the ChIP samples. The bamCompare function from Deeptools was used to normalize ChIP samples to Input or H3. log2 ratio (ChIP/ (Input or H3)) bigwig files were then used to generate metaplots and heatmaps across the different features in the genome (protein coding genes, Transposable elements and TEgenes (https://www.arabidopsis.org/download/index-auto.jsp?dir=%2Fdownload_files%2FGenes%2FTAIR10_genome_release%2FTAIR10_gff3)).

### Defining chromatin states

Aligned ChIP-seq BAM files for 27 chromatin features were generated as described above. Aligned BAM files were then converted to BED format using .These genome wide BED files were then used for learning chromatin state by a multivariate Hidden Markov Model. We used the LearnModel program from ChromHMM (Ernst and Kellis 2017) with default settings and window size 200bp. We generated chromatin states from n=2 to n=50(n, number of chromatin states). We chose to analyze the 26 chromatin state model in depth.We analyzed multiple states models within a window of n=13 to n=27 states. We looked at states association with genomic features. We observed that increasing state numbers from 26 to 27 gave rise to biologically redundant states. Hence we chose to analyze 26 state models in depth.

### Analysis of TE groups

To identify Transposable elements that were associated with each chromatin state. TEs with length greater than 500 bp were considered for further analysis. Using Deeptools we performed hierarchical clustering on TEs. The State region file was used to create a signal bigwig file. Presence of state was assigned score “1” and absence of state was assigned “0”. This scored signal file for each state was then used in Deeptools to generate TE groups based on state enrichment. Not all the states overlapped with TEs above 500bp, Hence only heterochromatic states H1-H6 and I1 and F1 were used. lustering for TE groups was performed using n= 2 to n=10 clusters. Clusters with similar profiles were merged and we chose to analyze 7 unique groups of TEs.

### Analysis of chromatin landscape across protein coding genes

To identify Chromatin landscapes across protein coding genes. Using Deeptools we performed hierarchical clustering on all protein coding genes annotated in TAIR10. The State region file was used to create a signal bigwig file. Presence of state was assigned score “1” and absence of state was assigned “0”. This scored signal file for each state was then used in Deeptools to generate CL groups based on state enrichment across these genes. Clustering for protein coding genes was performed using n=2 to n=26 clusters. Clusters with similar profiles were merged and we chose to analyze 10 unique groups of genes each with a specific chromatin landscape.

### Analysis of gene quantiles in each chromatin landscape

For each euchromatic chromatin landscape (CL7-10), genes in each CL were divided in four quantiles (q1=lowest expression and q4=highest expression) based on the expression in TPM (Transcript per million). Deeptools v3.1.2(Ramírez et al. 2016) was used to generate heatmaps and average profiles for various ChIP-seq, ATAC-seq, Pol-II phosphorylation forms as well as CG methylation profiles across these gene quantiles(q1-q4) in each CL.

### Data availability

The genome-wide sequencing (ChIP-seq) generated for this study as well as published datasets (ChIP-seq, RNA-seq) utilized to support the findings in this study have been deposited on NCBI’s Gene Expression Omnibus (GEO) with the accession number XXXX.

Source data for all the main figures as well as supplementary figures have been provided with this manuscript. All other data supporting the conclusions of the study will be available from the corresponding author upon reasonable request.

### Code availability

All the code used to analyze the genome-wide sequencing data presented in this study, as described in the Methods are available upon request.

## Supplementary Figure Legends

**Fig. S1:**
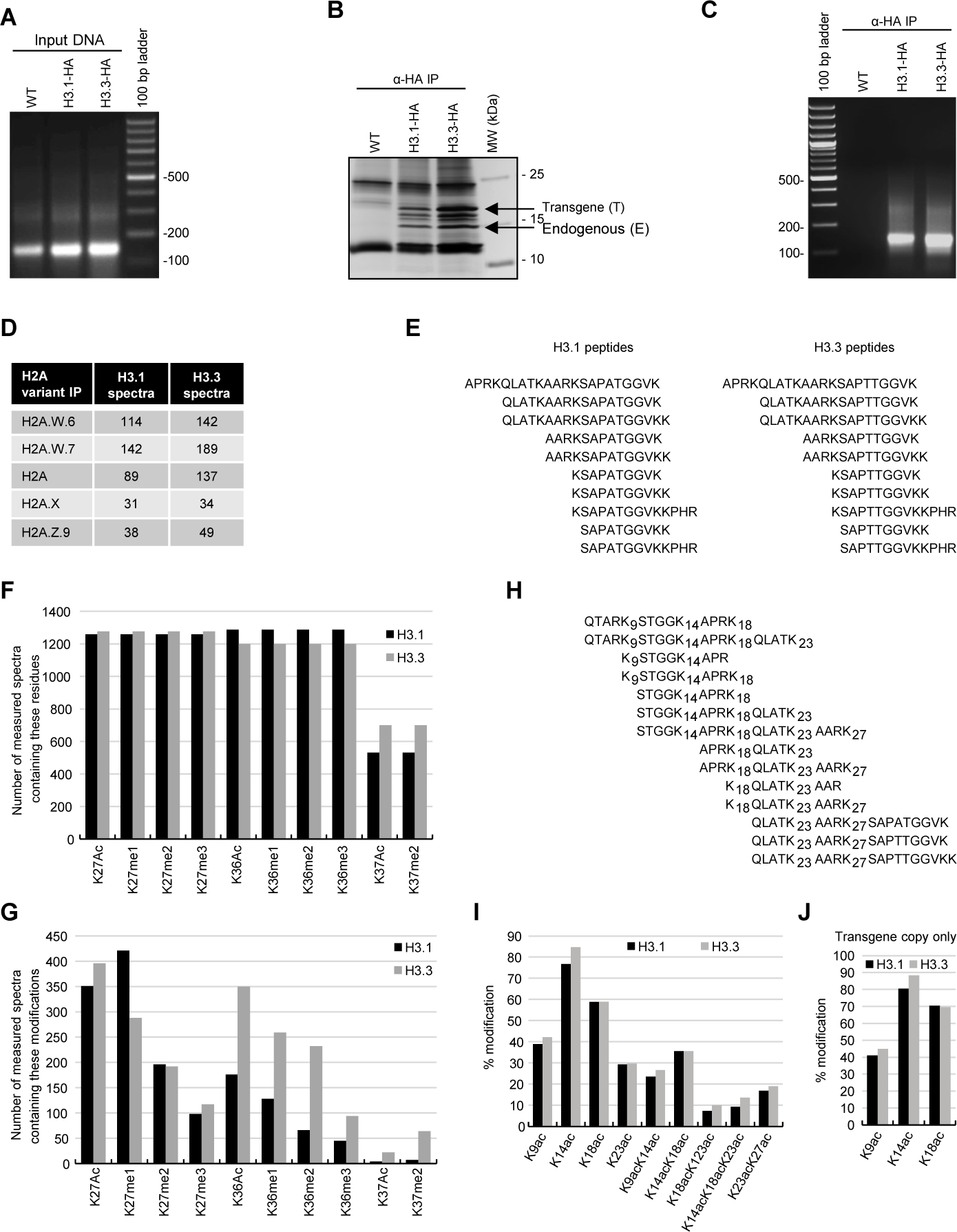
Biochemical analysis of the association between histone variants and histone marks. **(A)** DNA in input samples used for H3.1 and H3.3 immunoprecipitation. **(B)** Silver-stained gel of immunoprecipitated H3.1 and H3.3 mononucleosomes. **(C)** DNA extracted from immunoprecipitated H3.1 and H3.3 mononucleosomes. **(D)** Spectral counts of H3.1- and H3.3-specific peptides in immunoprecipitates with H2A variants. **(E)** H3.1- and H3.3-specific peptides used for the analysis of H3 K27 and K36 modifications. **(F)** Number of MS spectra measured that covered the indicated residues. **(G)** Number of measured MS spectra containing the indicated modifications. **(H)** Sequences of peptides used to evaluate H3 acetylation by MS. **(I)** Relative acetylation levels of the indicated H3 residues in H3.1 and H3.3 immunoprecipitates. **(J)** Relative acetylation levels of the indicated H3 residues on H3.1 and H3.3 revealed from transgenic copies of the respective H3 variants. Histone H3.1 and H3.3 form homotypic and heterotypic nucleosomes. Spectral counts of H3.1- and H3.3-specific peptides in the respective immunoprecipitates (T – transgenic, E – endogenous H31. and H3.3). (E) Number of MS spectra measured covering the indicated residues.

**Fig. S2:**
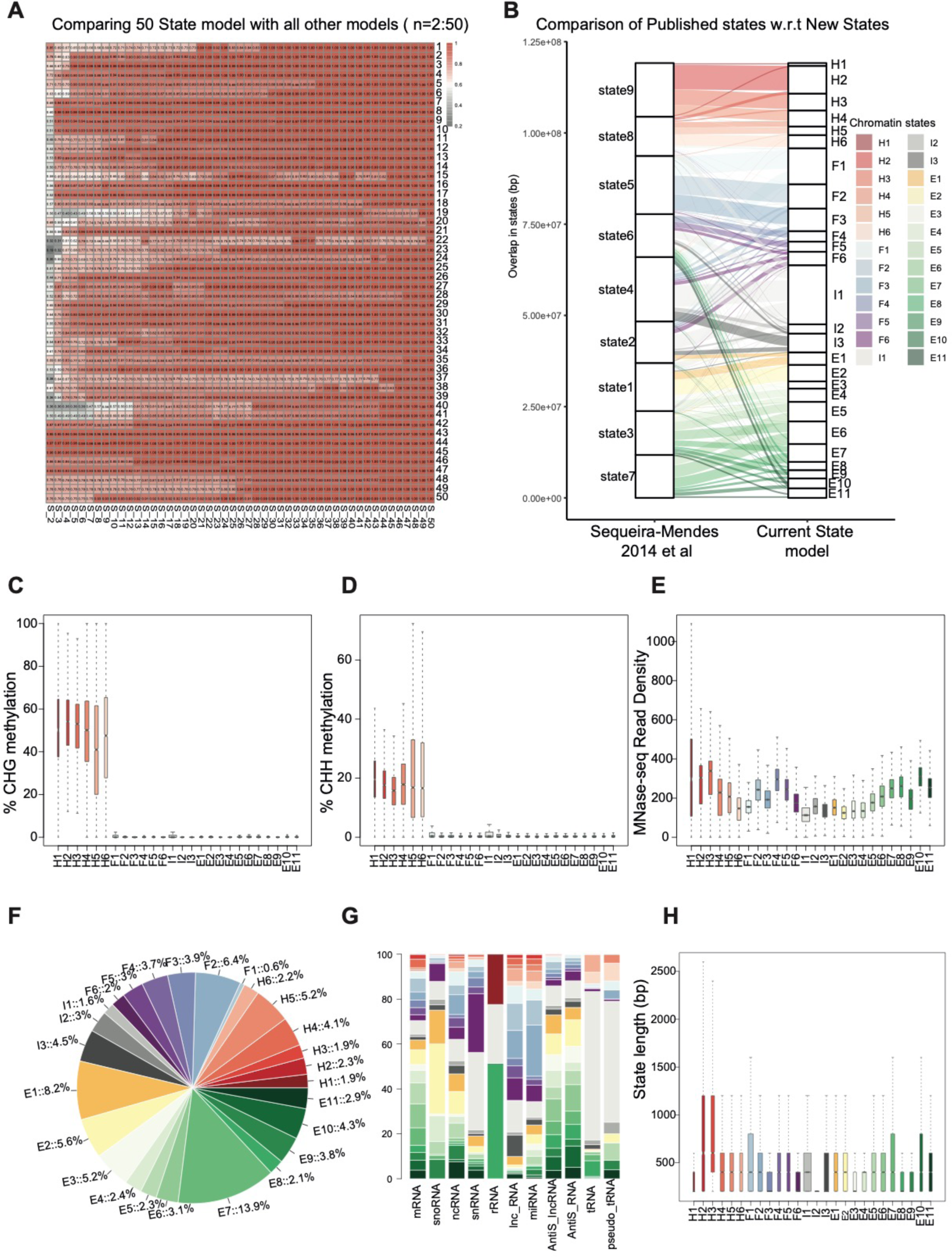
Comparison between published chromatin states and chromatin states and properties associated with states identified in the current study. **(A)** Heatmap showing the correlation across states from different numbers of chromatin states. States model with 50 states was used to calculate correlation with other models with lower numbers of states (n=49: n=2). **(B)** Flow diagram showing the overlap (in bp) between published states (Sequeria-Mendes 2014 et al) and the current state model. Color code for the flow diagram represents the chromatin states in the current model. **(C, D)** Boxplots showing methylation levels for CHG and CHH across states. **(E)** Boxplot showing nucleosome occupancy (MNase-seq read density) across different states. **(F)** Pie plot showing the genome % of each state. **(G)** Overlap between non-protein-coding gene features and states.

**Fig. S3:**
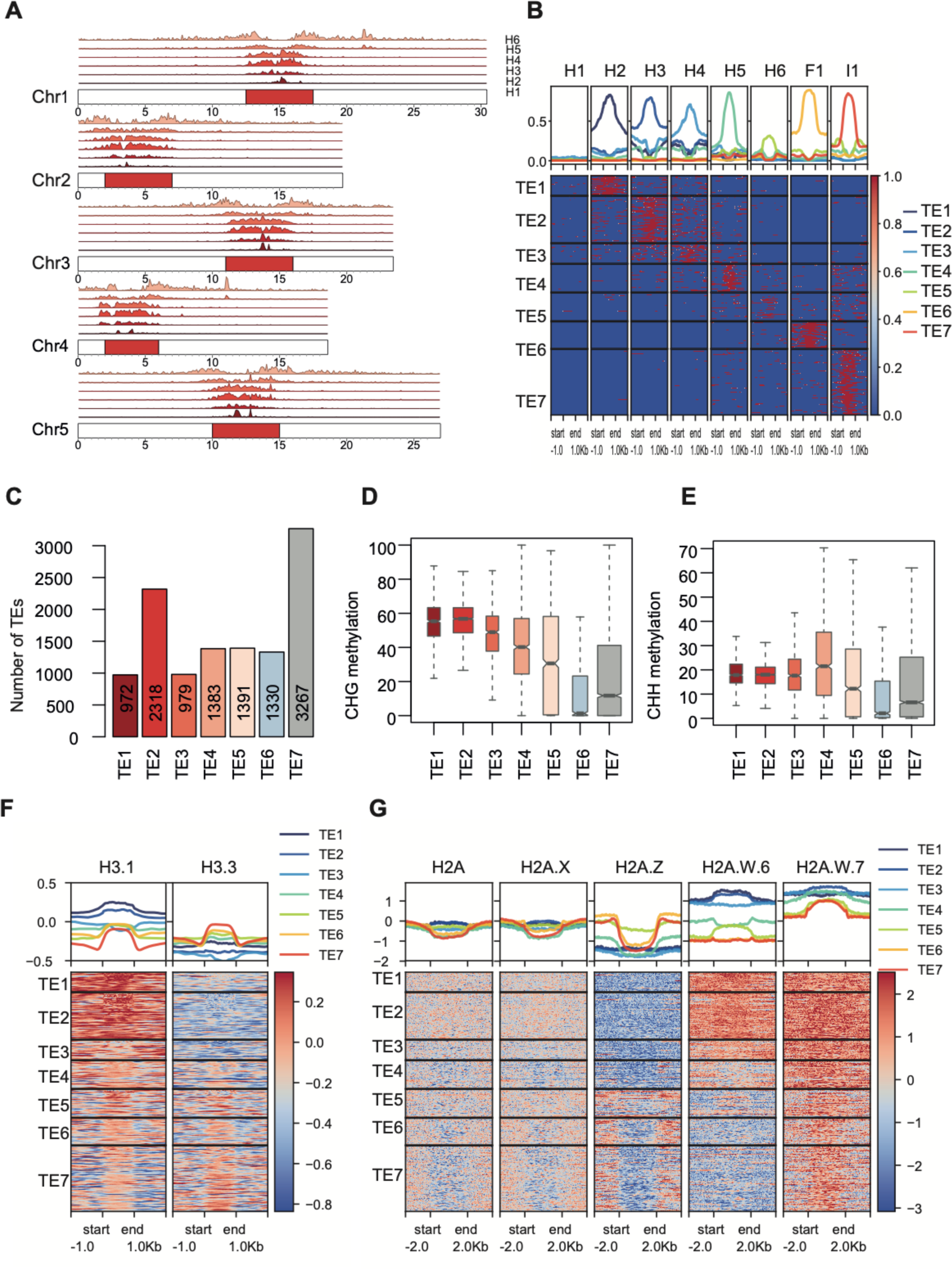
Transposable elements clusters and their properties. **(A)** Genomic distribution of heterochromatic states (H1–H6) [bottom to top] along all chromosomes in *Arabidopsis thaliana*. **(B)** Heatmap showing the occupancy of heterochromatic chromatin states (H1–H6) across TE groups TE1–TE7. **(C)** Bar plot showing the number of TEs in each TE group. **(D, E)** Boxplot showing the methylation levels across TE groups from left to right (CHG and CHH, respectively). **(F)** Aggregate ChIP-seq profiles and heatmaps for H3 variants across the TE groups. **(G)** Aggregate ChIP-seq profiles and heatmaps for H2A variants across the TE groups.

**Fig. S4:**
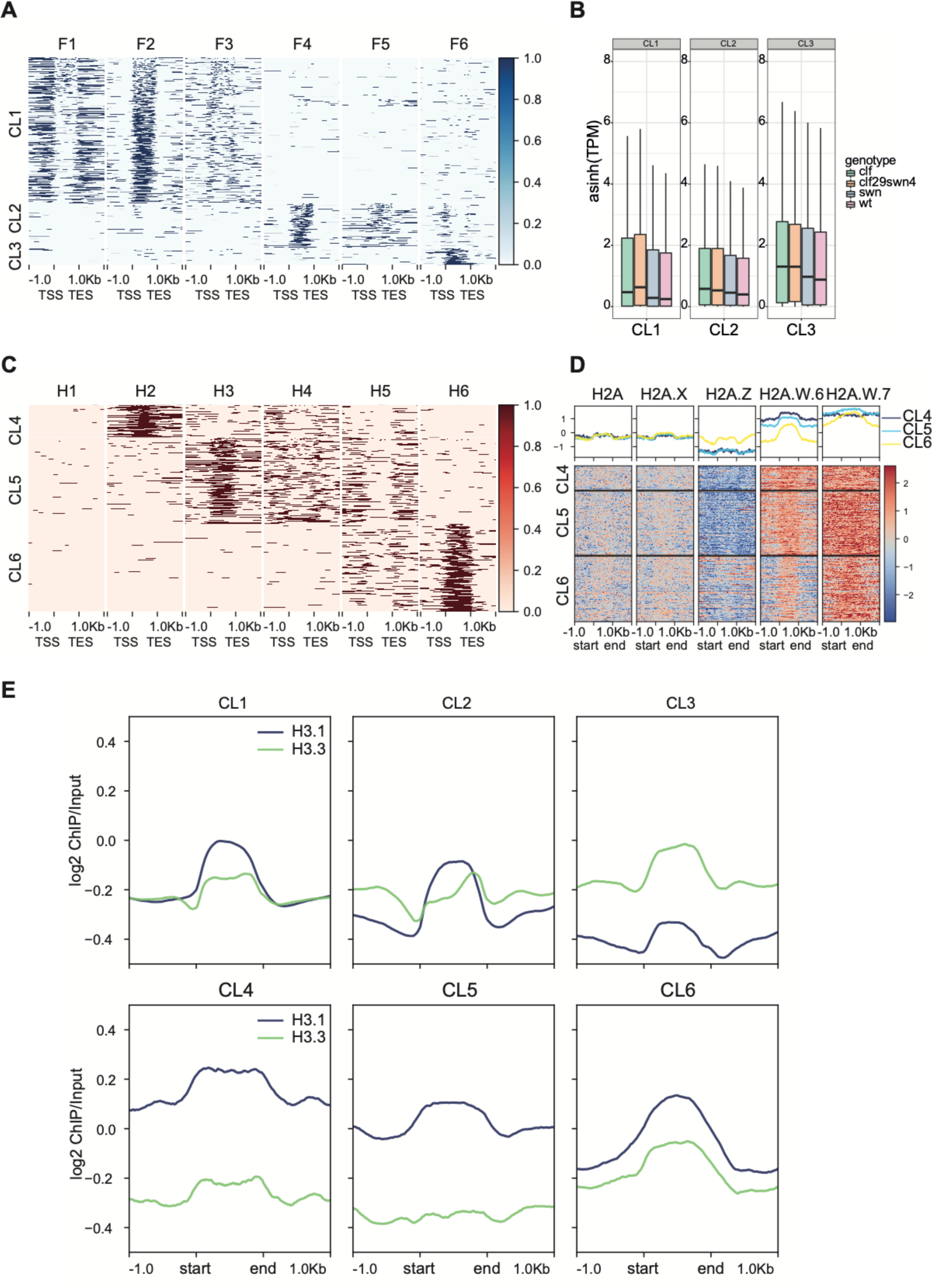
Chromatin landscapes across developmentally repressed genes. **(A)** Heatmap displaying the occupancy of facultative chromatin states (F1–F6) across the CL1–CL3 genes. **(B)** Boxplot showing the expression levels of CL1–CL3 genes in PRC mutants. **(C)** Heatmap displaying the occupancy of heterochromatic chromatin states (H1–H6) across gene clusters CL4–CL6. **(D)** Aggregate profile plots and heatmaps showing ChIP-seq signals for H2A variants across CL4–CL6. **(E)** Aggregate profile plots showing ChIP-seq signals for H3.1 and H3.3 along repressed genes (CL1–CL6).

**Fig. S5:**
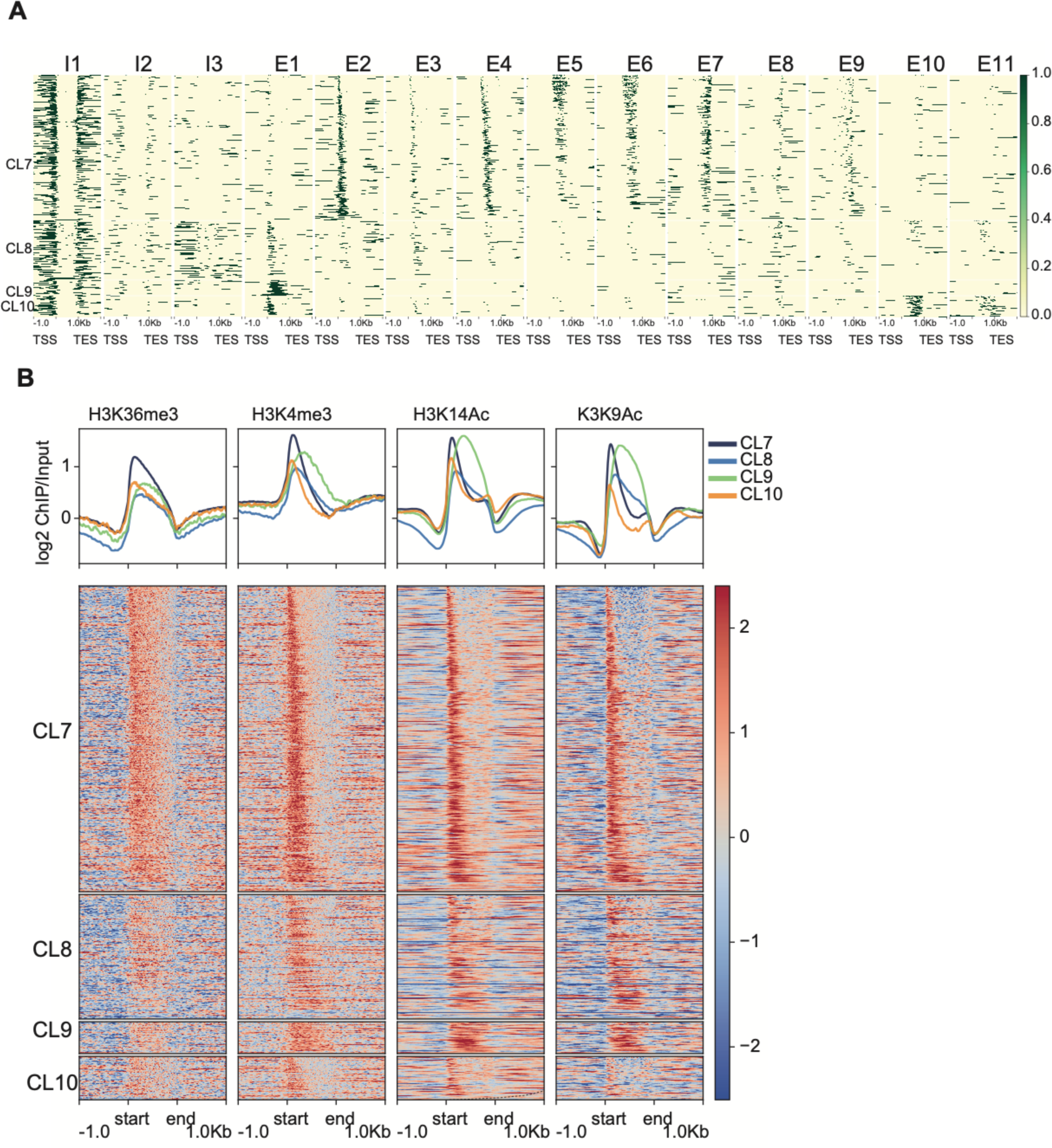
Expressed genes chromatin landscapes. **(A)** Heatmap showing the intergenic and euchromatic states across active protein-coding gene clusters (CL7–CL10). **(B)** Heatmap showing log2(ChIP/Input) for active chromatin marks across four types of gene clusters.

**Fig. S6:**
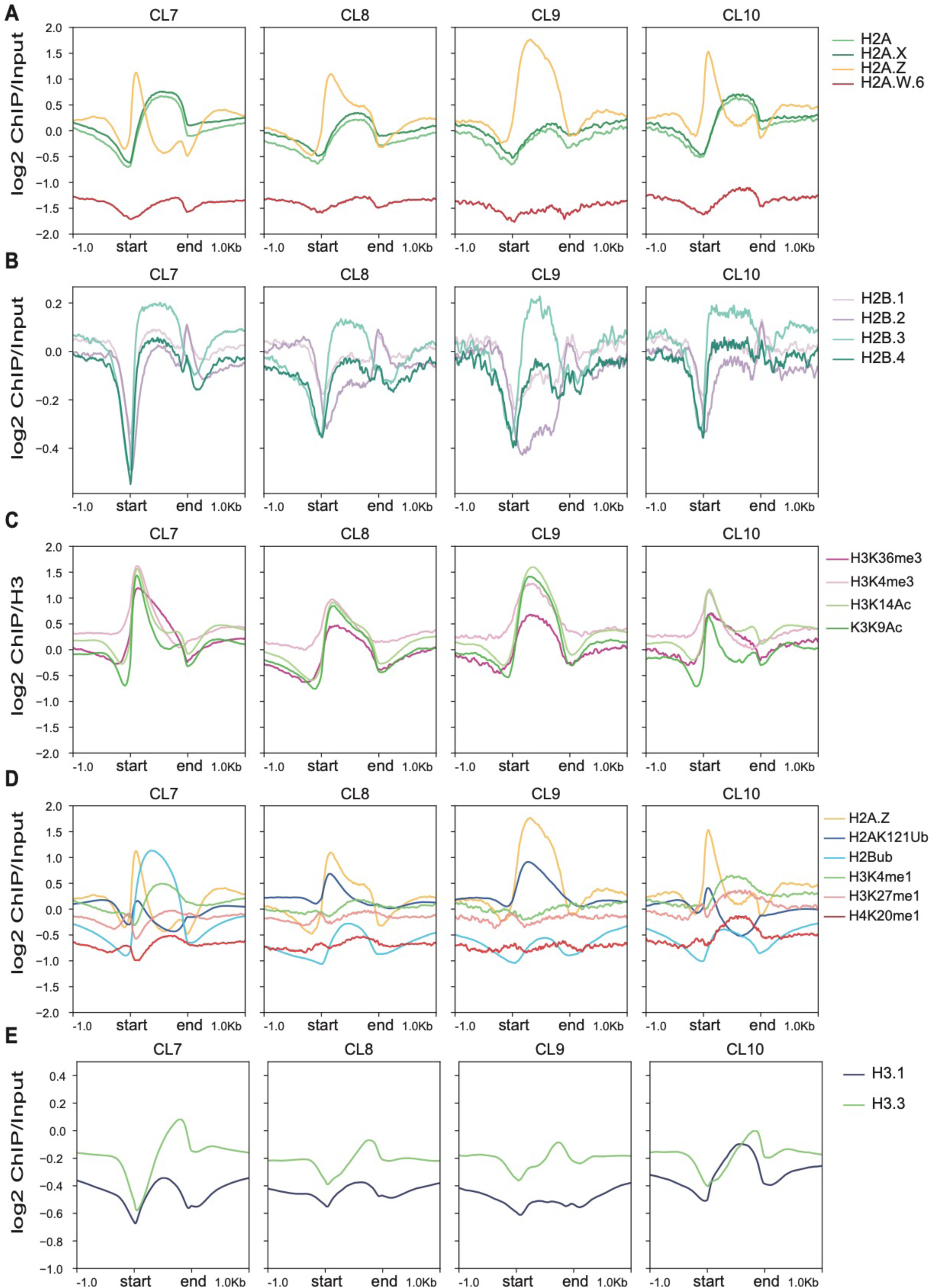
Expressed genes chromatin landscapes. Aggregation profiles across CL7–CL10 genes showing log2 (ChIP/Input) for chromatin marks. **(A)** H2A variants **(B)** H2B variants **(C, D)** Histone PTMs, and **(E)** H3 variants.

**Fig. S7:**
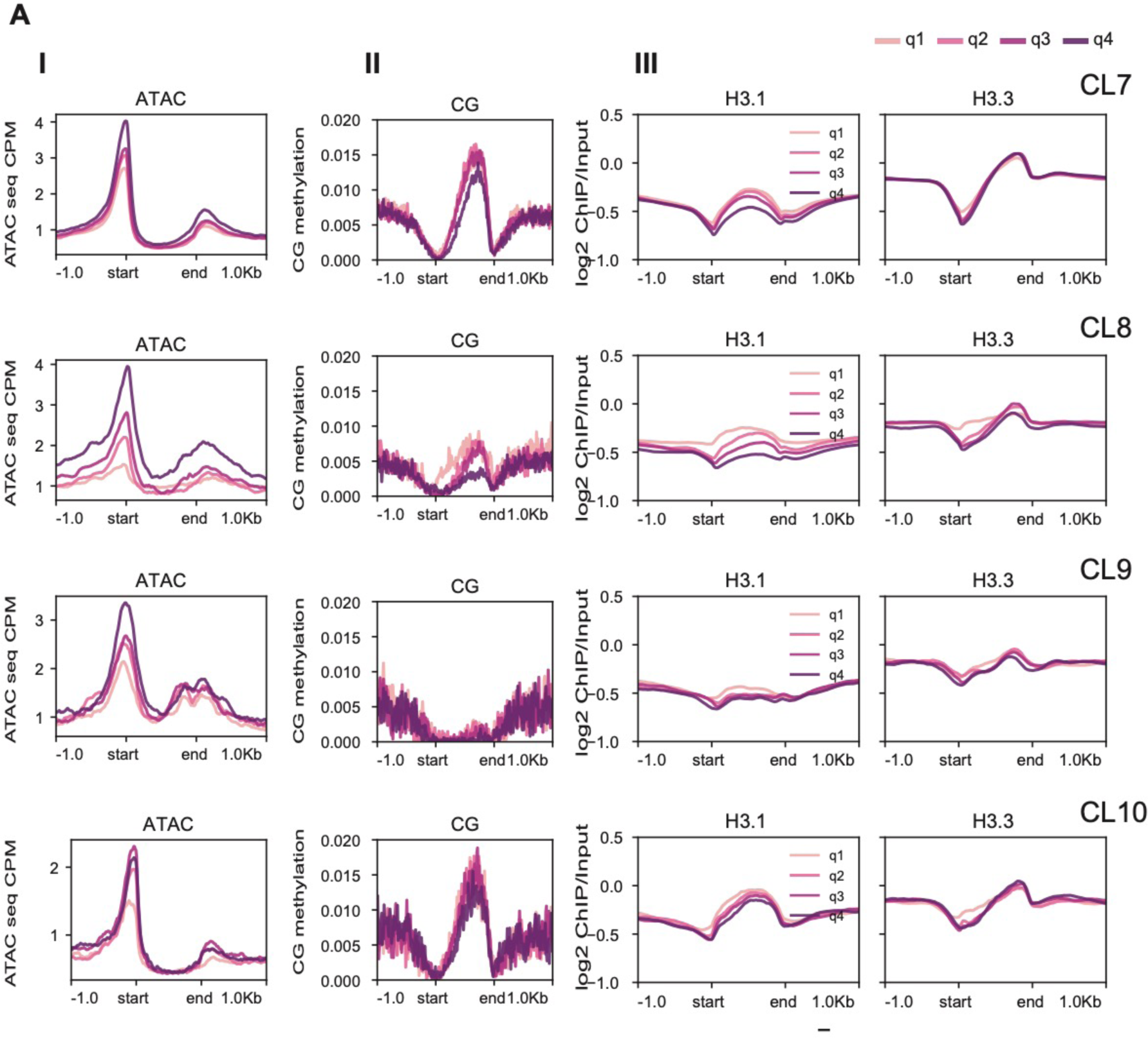
Expression quantiles within chromatin landscapes. **(A)** Aggregation profiles across expression quantiles (q1= lowest; q4=highest) in CL7–CL10**: I)** Accessibility (ATAC-seq signal) across gene quantiles**, II)** CG methylation levels, and **III)** log2 (ChIP/Input) signal for H3 variants across gene quantiles in each CL type.

**Fig. S8:**
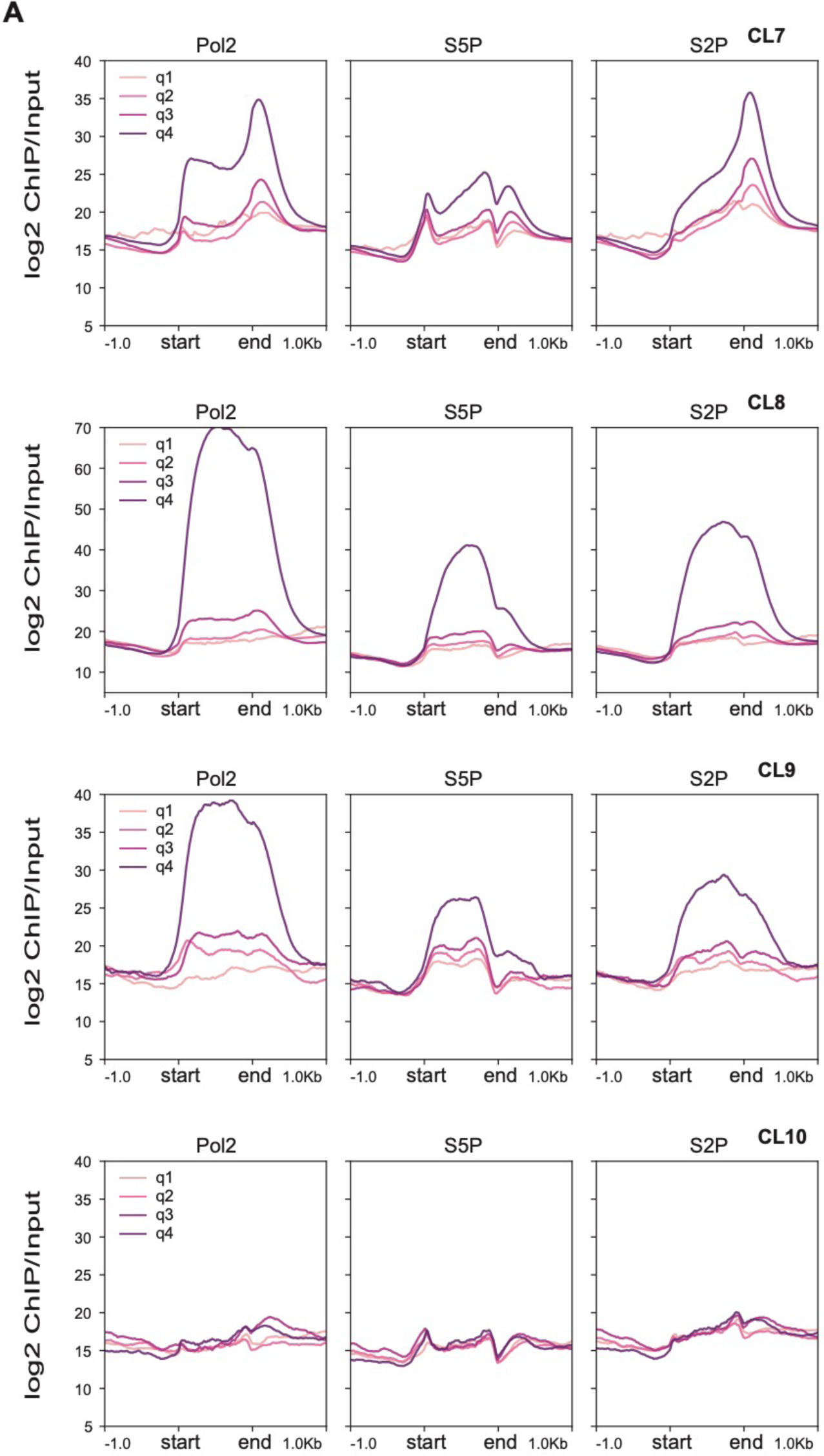
Expression quantiles within CLs and Pol II. **(A)** Aggregation profiles for Pol II occupancy across expression quantiles (q1= lowest; q4=highest) in each CL type (CL7– CL10).

**Fig. S9:**
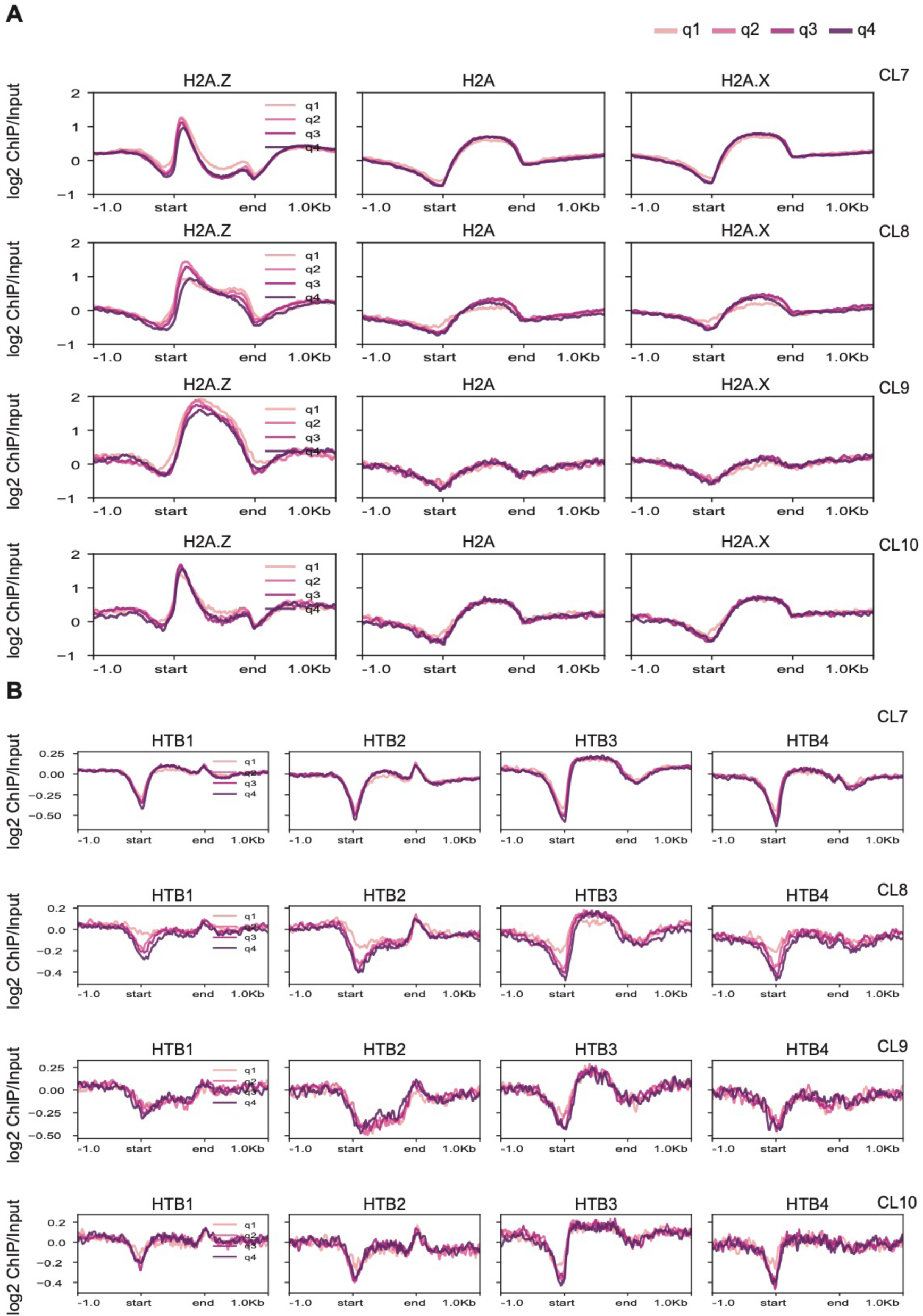
Expression quantiles within CLs and H2A and H2B variants. Aggregation profiles for log2 ChIP/Input signal for **(A)** H2A variants and **(B)** H2B variants across expression quantiles (q1= lowest; q4=highest) in each CL type (CL7–CL10).

**Fig. S10:**
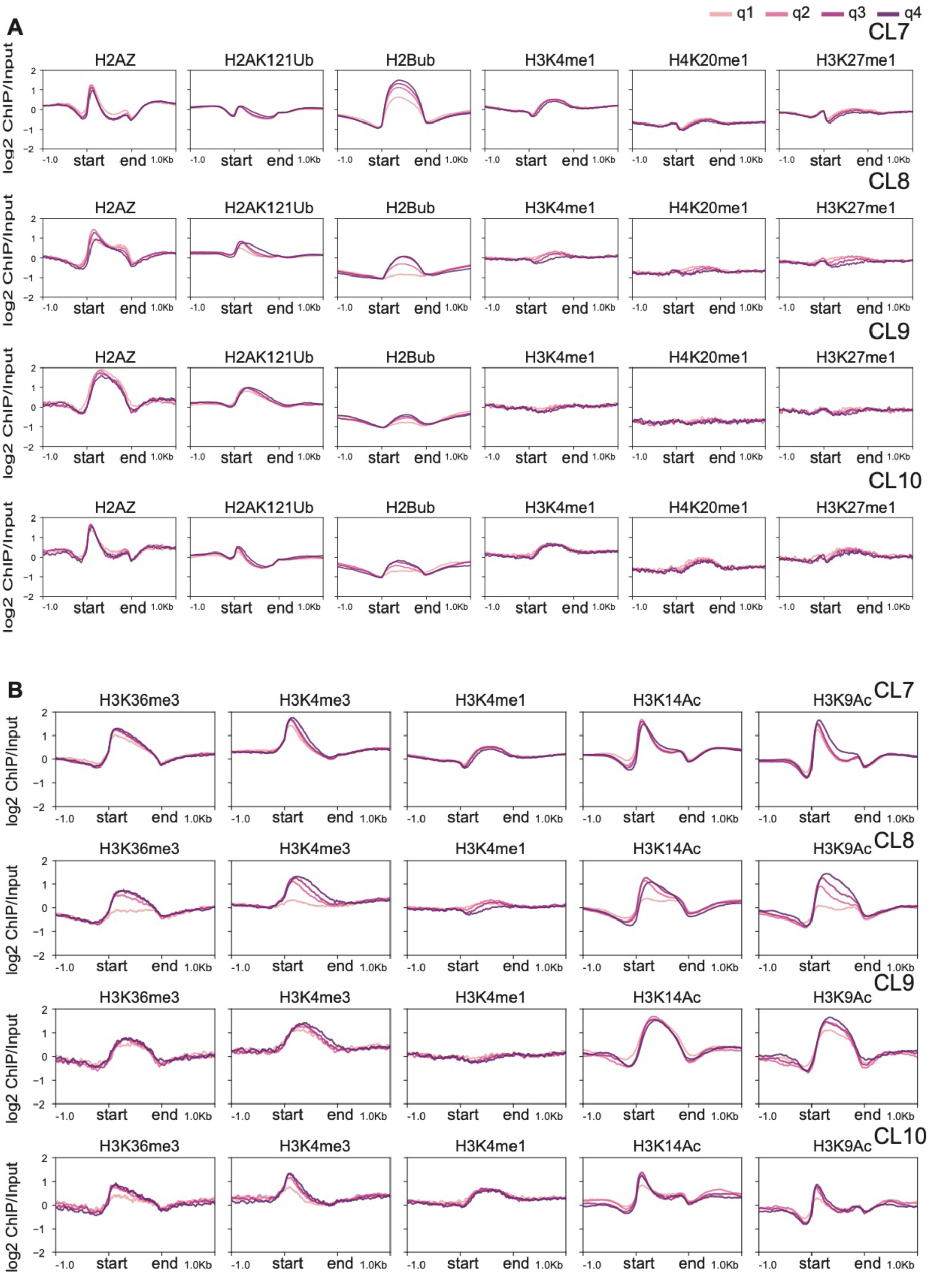
Expression quantiles within CLs and H3 modifications. Aggregation profiles for log2 ChIP/Input signal for **(A)** H2AZ and multiple H3 modifications across expression quantiles (q1= lowest; q4=highest) in each CL type (CL7–CL10).

